# Adhesion-derived condensates control component availability to regulate adhesion dynamics

**DOI:** 10.1101/2025.05.08.652869

**Authors:** Michal Dibus, Giray Enkavi, Megan Chastney, Ilpo Vattulainen, Johanna Ivaska

## Abstract

Integrin adhesion complexes mediate cell-extracellular matrix (ECM) interactions and undergo dynamic remodelling to regulate cell adhesion and migration. Here, we demonstrate that tensin 1 (TNS1), a multidomain adhesion adaptor protein linking active integrins with the actin cytoskeleton, undergoes phase separation in cells. Endogenous TNS1 condensates are formed upon focal adhesion disassembly or limited integrin–ECM engagement in both 2D and 3D environments, acting as reservoirs for inactive adhesion proteins. Combining functional experimental approaches with phosphoproteomics, we identify the TNS1 intrinsically disordered region as the main driver of TNS1 condensation and demonstrate a negative regulatory role of phosphorylation on condensate assembly upon activation of stress-responsive kinases. Finally, we confirm the functional effects of phosphorylation-dependent TNS1 condensation on adhesion dynamics and cell migration. Together, our findings highlight TNS1 condensation as a regulatory mechanism controlling local availability of inactive adhesion proteins, with direct implications on adhesion dynamics and cell behaviour.

## Introduction

Integrin adhesion complexes mediate cell interactions with extracellular matrix (ECM) components and are crucial in regulating various cellular processes, including cell survival and migration. At the onset of adhesion, small and short-lived nascent adhesions are formed that can further mature into focal adhesions (FAs) as a result of changes in protein conformation, molecular composition, and actomyosin-mediated contractility^1,2^. Since FAs represent dynamic multi-protein assemblies^3^, rapid recruitment of individual FA components must be tightly controlled. While the nanoscale molecular composition of FAs is relatively well understood^4^, mechanisms governing FA component availability to facilitate their rapid recruitment at adhesion sites remain elusive. In addition, there is increasing evidence suggesting liquid-like FA organisation^5,6^, prompting further studies into how liquid-like properties of individual FA components influence their regulation and dynamics^7^.

Liquid–liquid phase separation is a biophysical process in which biomolecules spontaneously demix from solution to form condensates, whose properties are shaped by the surrounding physicochemical environment^8^. It is well-established that (nano-scale) phase separation drives macromolecule compartmentalisation in cells, resulting in biogenesis of membraneless organelles, such as processing bodies (P-bodies), stress granules and nuclear bodies^9^. In addition, recent studies have highlighted the role of biomolecular condensates in regulating diverse array of signalling events and cellular processes across the kingdoms of life, revolutionizing the way we understand them. For instance, condensation of the tight junction protein ZO-1 at cell membrane interfaces has been proved essential for the assembly of tight-junction belts in polarized epithelial tissues^6,10,11^. Similarly, phase separation of the WNK1 kinase has been shown to play a crucial role in cellular response to hyperosmotic stress^12^. Finally, in plants, FLOE1 condensates modulate seed germination in response to unfavourable environmental conditions^13^.

A number of recent studies reported key nascent and FA components to form biomolecular condensates *in vitro* and in cells, regulating their localisation and function^5,14–19^. However, while these works provide valuable insights into FA-assembly mechanisms, many conclusions are drawn from *in vitro* reconstitutions of purified proteins, omitting the complexity of the cellular environment. The large multidomain tensin family members (tensin 1-3; TNS1-3) are key components of adhesion complexes and exert their functions by linking the actin cytoskeleton with the ECM through direct and indirect interactions with integrin heterodimers ^20,21^. The shorter, oncogenic tensin 4 acts as a promigratory protein supporting growth-factor-receptor-mediated malignant processes^22,23^. Work from us and others has implicated TNS1 in maintaining active integrin pools and adhesion reinforcement^24–27^ and facilitating tension-dependent FA maturation into centrally localized fibrillar adhesions^28–31^. Furthermore, the importance of TNS1 dynamics in regulating integrin adhesions is exemplified in vascular endotoxemia, where rapid recruitment of TNS1 reinforces FAs, ultimately leading to increased vascular leakage^32^. Although key roles of TNS1 in FA regulation have been demonstrated across species^20,24^, the mechanisms underlying TNS1 dynamics in these processes remain poorly understood. In this work, we provide evidence that TNS1 forms functional, endogenous protein condensates in a cellular context, and, as such, TNS1 selectively microcompartmentalises inactive adhesion components to functionally relevant cytoplasmic reservoirs. Using a combination of experimental approaches and molecular simulations, we identified regions critical for driving TNS1 condensation and established their significance in regulating TNS1-mediated biological functions. Finally, we demonstrated the regulatory role of TNS1 phosphorylation in condensate disassembly and control of FA dynamics.

## Results

### TNS1 forms biomolecular condensates in cells

In our previous studies focusing on the role of TNS1 in metabolism-mediated regulation of integrin activity^28^, we noted the formation of TNS1 biomolecular condensates similar to those observed in phase-separated proteins. To characterise these condensates and investigate whether TNS1 can undergo (nano-scale) phase separation, we generated a U2OS (human osteosarcoma) cell line with inducible GFP-TNS1 expression. Analysis of individual cells with varying TNS1 expression levels revealed a strong positive correlation between TNS1 expression and the total TNS1 condensate area in individual cells (Figures 1A and 1B), suggesting TNS1 condensation is concentration-dependent. We next focused on the dynamics of GFP-TNS1 condensates in living cells and confirmed that they are able to both fuse (Figure 1C, Movie S1) and split into smaller ones (Figure 1D, Movie S2). We also observed TNS1 condensate formation during FA disassembly (Figure S1A, Movie S3). While condensates proximal to FAs exhibited very limited mobility, more distal condensates exhibited dynamic movement in the cytosol (Figures S1B – S1D, Movie S4). Using fluorescence recovery after photobleaching (FRAP) we also demonstrated rapid protein turnover (ca. 49 % recovery, half-life 27.58 s ± 1.01 s) between the GFP-TNS1 condensates and the soluble pool of the protein (Figures 1E and 1F, Movie S5). Finally, one of the major features of biomolecular condensates is the absence of membranes around these structures. To evaluate this, we first stained the GFP-TNS1-expressing cells with fluorescently-conjugated wheat germ agglutinin (WGA) that binds to glycoproteins on the nuclear, endosomal and plasma membranes. WGA staining showed no overlap with GFP-TNS1 condensates confirming that they are not a part of the endosomal compartment (Figure 1G). However, given the limitations of WGA staining, we further employed correlative light electron microscopy (CLEM) to confirm the absence of membrane around TNS1 condensates. Electron micrographs correlated with immunofluorescence (IF) images revealed the absence of any discernible membrane around the GFP-TNS1, in contrast to the adjacent membrane-limited organelles, such as mitochondria, endoplasmic reticulum or vesicles (Figure 1H, annotated micrograph in Figure S1E). Moreover, the condensates appeared as electron-dense granules similar to other phase-separated cellular structures, such as Cajal bodies^33^, P-bodies^34^ and stress-granules (SG)^35,36^. Taken together, in line with the accepted guidelines^37^, we provide strong evidence that TNS1 condensates are dynamic membraneless particles able to exchange material with the surrounding cytoplasm, confirming that TNS1 can undergo phase separation in a concentration-dependent manner *in cellulo*.

**Figure 1.**
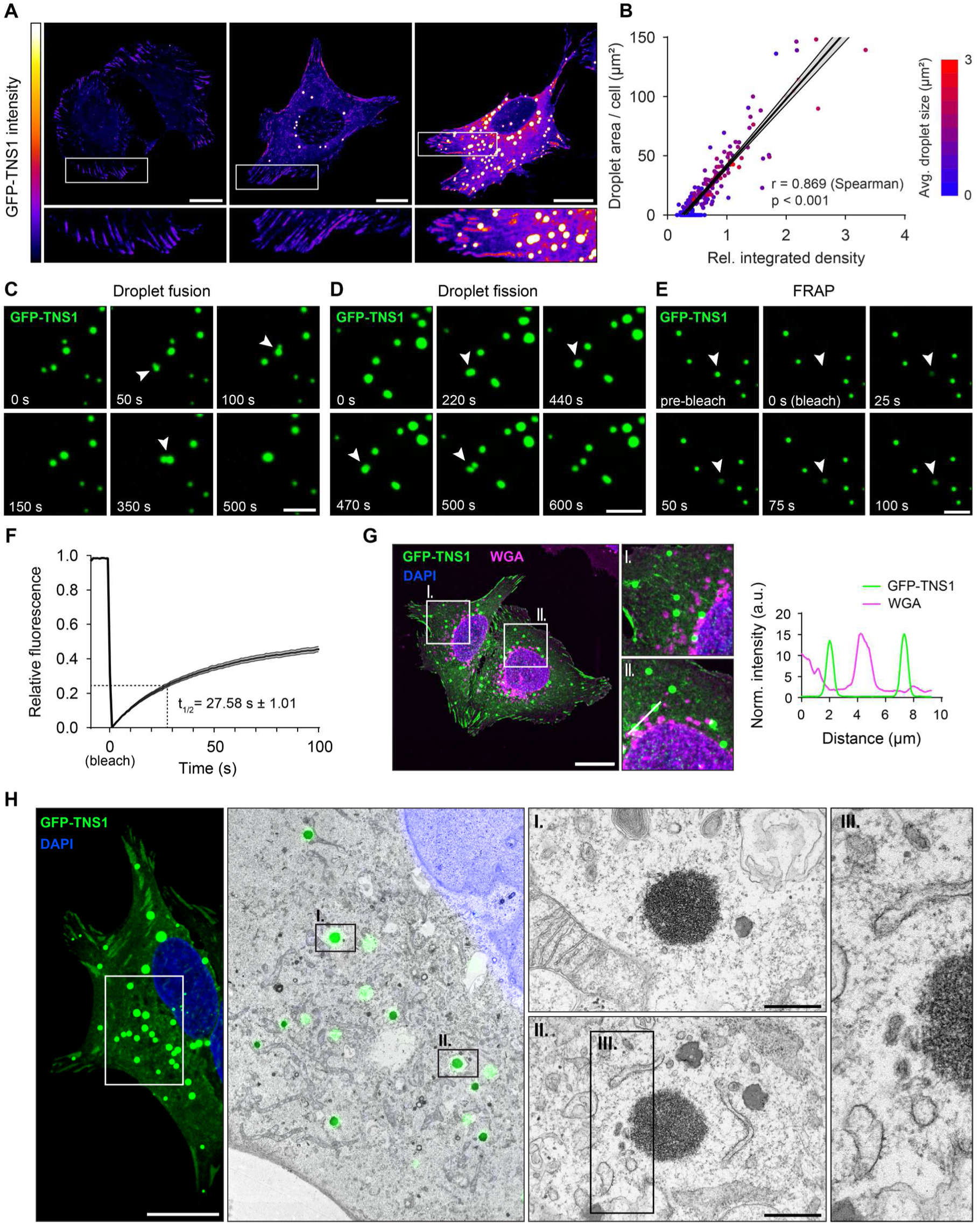
TNS1 undergoes phase separation in a concentration-dependent manner. A) Representative confocal images of U2OS cells with varying expression of GFP-TNS1. Scale bar 20 μm. B) Correlation between relative expression of GFP-TNS1 in U2OS cells and the total condensate area per cell, represented as linear regression with 95% confidence intervals. Color-coded data points represent average condensate size in each of the quantified cells. Quantified from n = 189 cells. C) Representative confocal time-lapse images of GFP-TNS1 condensate fusion (indicated by arrowheads) in U2OS cells. Cells were imaged for 10 min at 10 s intervals. Scale bar 5 μm. D) Representative confocal time-lapse images of GFP-TNS1 condensate relaxation (indicated by arrowheads) in U2OS cells. Cells were imaged for 10 min at 10 s intervals. Scale bar 5 μm. E) Representative images from FRAP experiments. After bleaching one GFP-TNS1 condensate per cell, cells were imaged for 100 s at 1 s intervals. Bleached condensate is indicated by arrowhead. Scale bar 5 μm. F) Average FRAP curve from measurement of n = 34 condensates plotted with standard error of mean (SEM). Half-life of maximum recovery (t_½_) with SEM is indicated. G) Representative confocal images of U2OS cells expressing GFP-TNS1, stained with fluorescently-conjugates wheat germ agglutinin (WGA) and DAPI. Intensity profile plot depicts relative GFP-TNS1 and WGA intensities along the white line indicated in closeup II. Scale bar 20 μm. H) Representative images from correlative light-electron microscopy of GFP-TNS1 condensates in U2OS cells. Scale bar on confocal image 20 μm, scale bars on electron micrographs 500 nm.

### Endogenous TNS1 condensation is controlled by actomyosin contractility and ECM ligand availability in 2D and 3D

To study TNS1 condensation at endogenous protein levels, we generated a U2OS cell line with endogenous TNS1 tagged with the bright monomeric green fluorescent protein, mGreenLantern (mGL)^38^ (Figures S2A – S2D). To examine the response of TNS1-containing FAs to different ECM stimuli, we seeded these cells on glass coverslips coated either with various integrin-binding ECM ligands or with poly-L-Lysine (PLL) to mimic conditions with low ligand availability. We observed no significant difference in FA area between cells seeded on fibronectin (FN), laminin (LMN), collagen IV (COL) and vitronectin (VTN), however, FA area was significantly reduced in cells seeded on PLL (Figures 2A and 2B). Interestingly, this correlated with a substantial increase in the amount of small, circular mGL-positive spots likely corresponding to TNS1 condensates and nascent adhesions (Figures 2C and 2D). To verify whether some of these spots represent TNS1 condensates, we generated orthogonal projections of cells seeded either on FN or PLL. While in cells seeded on FN all endogenous TNS1 was localized to the basal plane corresponding to FAs, in cells seeded on PLL several mGL-TNS1 spots were observed in higher cell planes (Figure 2E), indicating TNS1 forms condensates at endogenous levels in response to limited integrin-ECM engagement. To extend our observations further to a 3D environment, we embedded the cells into a 3D fibrillar collagen matrix. After 3 h, the cells had already formed prominent protrusions with TNS1 adhesions anchored on the underlying collagen fibres (Figure 2F, left panel). Interestingly, in less protruding cells, we observed several cytoplasmic TNS1 condensates (Figure 2F, right panels), suggesting their potential role in the local buffering of TNS1 amounts. We then performed a small screen using inhibitors of signalling pathways with established roles in regulating FA dynamics and examined their effect on TNS1 condensate formation in cells with endogenously tagged TNS1. While most inhibitors targeting canonical pathways downstream of integrin receptor activation and protein stability did not have a significant effect on TNS1 condensate formation (Figures 2G and S2E), treatment with compounds affecting actomyosin contractility and FA stability (ROCK inhibitors H1152, Y27632; Myosin-II inhibitor Blebbistatin; CDK1 inhibitor Ro3306) led to a markedly increased amount of TNS1 condensates, mostly as a result of FA disassembly (Figures 2G and 2H). Taken together, our results suggest that endogenous TNS1 condensates are formed in response to limited integrin ligand availability and decreased actomyosin contractility, and their formation could play a role in microcompartmentalisation and buffering of local protein concentration in both 2D and 3D.

**Figure 2.**
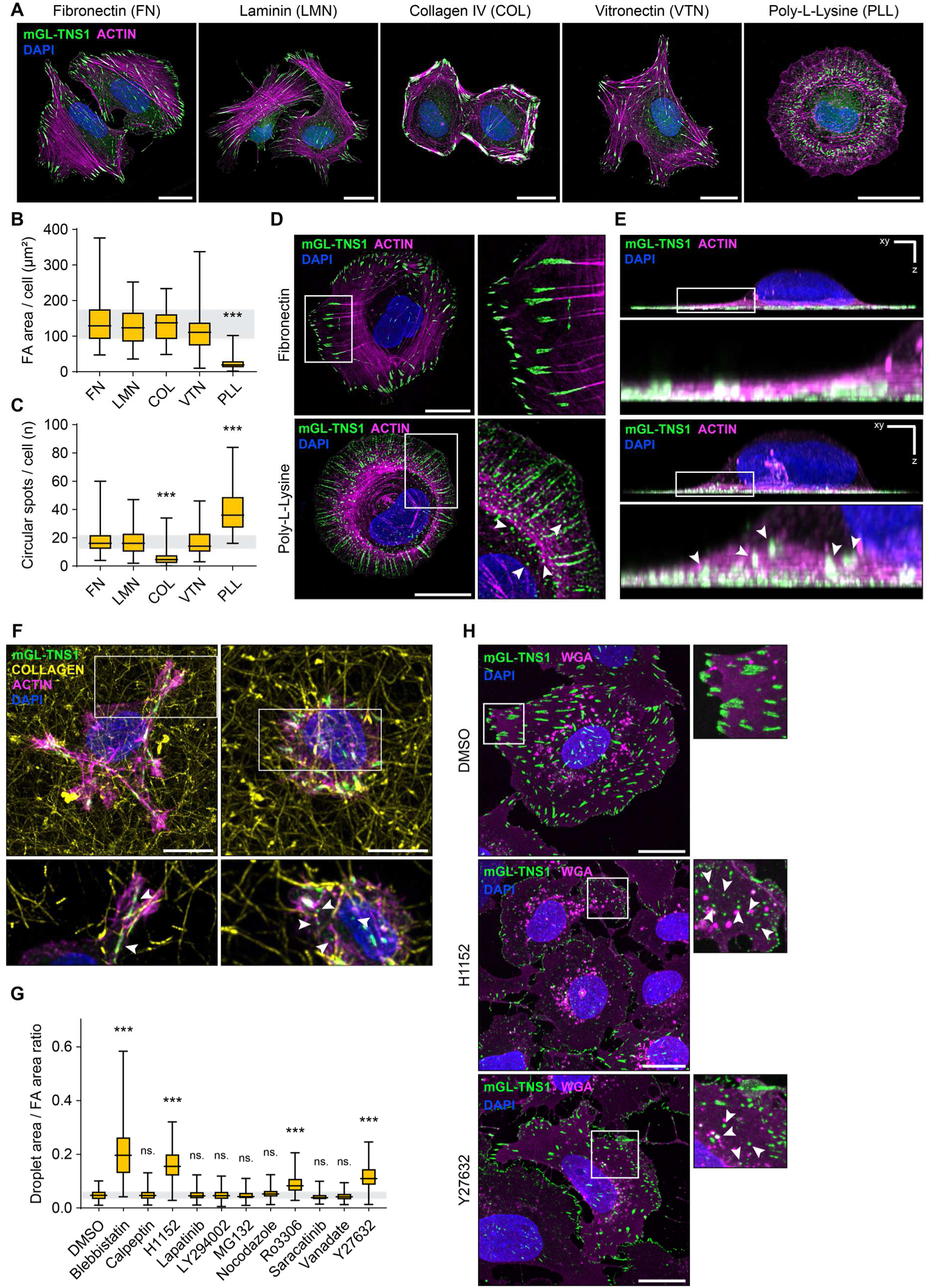
Endogenous TNS1 condensation is controlled by actomyosin contractility and ligand availability in 2D and 3D. A) Representative confocal images of U2OS cells with endogenously tagged TNS1 (mGreenLantern TNS1, mGL-TNS1) seeded on different integrin ligands or poly-L-Lysine as indicated. Images have been deconvolved using Huygens Professional (SVI). Scale bars 20 μm. B) Quantification of FA area per cell from n = 55-84 cells per condition. Statistical analysis was performed using Kruskal-Wallis test with Dunn’s multiple comparisons test. Indicated statistical significance level is valid for comparison with each of the other conditions. p < 0.001 (***). C) Quantification of number of circular spots per cell as in Figure 2B. Statistical analysis was performed using Kruskal-Wallis test with Dunn’s multiple comparisons test. Indicated statistical significance level is valid for comparison with each of the other conditions. p < 0.001 (***). D) Representative confocal images of U2OS cells with mGL-TNS seeded either on fibronectin or poly-L-Lysine and stained for F-Actin and DAPI. Some of the mGL-TNS1 condensates are indicated with arrows. Images have been deconvolved using Huygens Professional (SVI). Scale bar 20 μm. E) Orthogonal rendering of cells from Figure 5D. mGL-TNS1 condensates are indicated by arrows. Projection was performed in Huygens Professional (SVI). Scale bar 5 μm. F) Representative confocal images of U2OS mGL-TNS1 cells embedded in 3D fibrillar rat tail collagen conjugated with Atto647N. Cells were stained for F-Actin and DAPI. Maximum 3D projections are shown, closeups represent a maximum projection of 4 stacks at 0.72 μm. Arrows in the closeups indicate either FAs in contact with the underlying collagen fibres (left) or mGL-TNS1 condensates (right). Scale bars 10 μm. G) Representative confocal images of mGL-TNS1 U2OS cells treated with DMSO, H1152 or Y27632 inhibitors (both ROCK inhibitors). Cells were stained with WGA and DAPI. mGL-TNS1 condensates are indicated with arrows. Scale bars 20 μm. H) Quantification of the mGL-TNS1 droplet area and FA area ratio from n = 101-124 cells treated with the indicated compounds. Representative images are shown in Figures 2G and S2E. Statistical analysis was performed using Kruskal-Wallis test with Dunn’s multiple comparisons test against DMSO. p < 0.001 (***).

### TNS1 condensates act as a selective reservoir for inactive FA proteins

To study the molecular composition of TNS1 condensates and their potential role in buffering the concentration of FA components, we employed proximity biotinylation using BioID in U2OS cells with either low or high expression of TNS1-Myc-BioID (bait). While in the low-expressing cells the bait predominantly localised to FAs, higher TNS1-Myc-BioID expression induced condensate formation allowing us to study the phase separation-dependent TNS1 proximity interactome (Figures 3A, S3A and S3B). Statistical analysis of the BioID proteomic data using SAINTexpress^39^ identified 160 high confidence hits across 4 independent replicates (Figure S3C). As expected, gene ontology (GO) analysis of identified proteins revealed a significant enrichment in GO cellular component terms related to canonical TNS1 functions in linking cell adhesions with the actin cytoskeleton (Figure 3B). To validate the results of the screen and to identify FA-associated proteins in the TNS1 condensates, we performed IF staining against a set of FA proteins in U2OS cells with inducible GFP-TNS1 expression. While tensin 3 (TNS3), zyxin (ZYX), paxillin (PXN) and p130-Crk associated substrate (CAS) exhibited strong recruitment to TNS1 condensates, other FA-associated proteins such as focal adhesion kinase (FAK), talin 1 (TLN1), talin 2 (TLN2), vinculin (VCL) or KN Motif And Ankyrin Repeat Domains 2 (KANK2) showed limited to no FA localisation (Figures 3C and 3D, uncropped representative images for each staining are shown in Figure S3F), suggesting that protein recruitment to TNS1 condensates is of a selective nature. We further assessed the canonical phosphorylation status of these proteins downstream of active integrin signalling by staining with a general phospho-tyrosine (pY) antibody or antibodies recognising phosphorylated CAS, PXN or FAK. While all these antibodies specifically recognised the respective phosphorylated proteins at FAs, the staining was completely absent from GFP-TNS1 condensates (Figures 3E and 3F, uncropped representative images for each staining are shown in S3G). To validate these results also at the endogenous level, we seeded the U2OS cells with endogenously tagged TNS1 (mGL-TNS1) on PLL and stained for either PXN or phosphorylated PXN. Indeed, while PXN did colocalise with the mGL-TNS1 spots, phosphorylated PXN localised predominantly in the belt of nascent adhesions at the cell periphery and not in TNS1 condensates (Figure 3G). Our results suggest a novel role for TNS1 condensates in buffering the local concentration of FA proteins, representing a mechanism to maintain inactive FA proteins readily available for rapid recruitment during FA assembly. To further support this notion, we examined whether an increase in TNS1 expression could acutely stabilize the levels of FA components by their recruitment to TNS1 condensates. To test this, we induced GFP-TNS1 expression in U2OS cells using doxycycline for either 12 or 24 h and compared the expression levels of the selected proteins to those in uninduced cells. An increase in GFP-TNS1 expression led to significantly elevated levels of ZYX and PXN – the FA components that are recruited into TNS1 condensates. In contrast, the levels of TLN1 and VCL, which do not localise to TNS1 condensates, were not affected by changes in GFP-TNS1 expression (Figure 3H and 3I). Taken together, our results suggest that TNS1 condensates act as a selective reservoir for inactive FA components and can acutely stabilize protein levels of recruited FA proteins.

**Figure 3.**
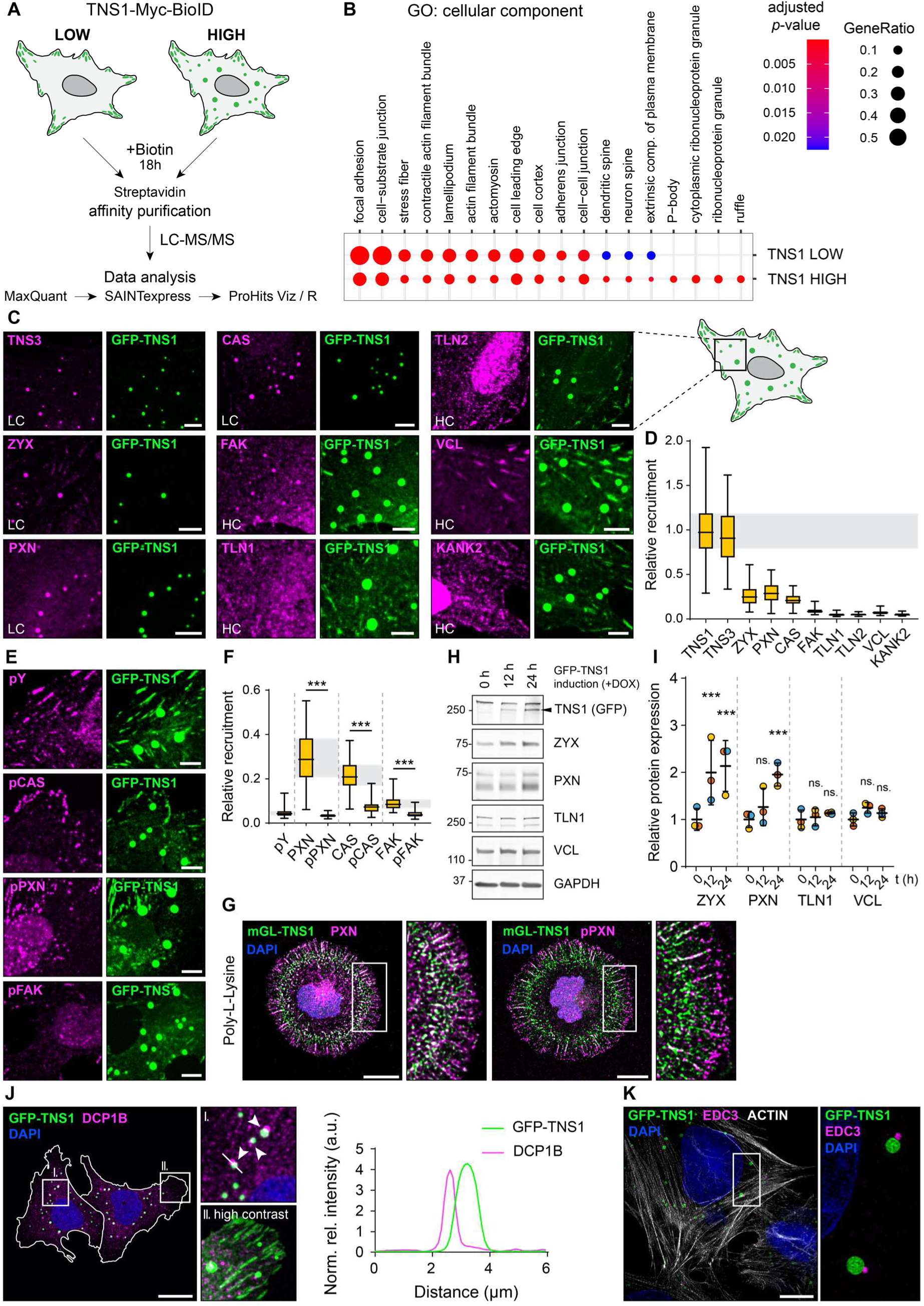
TNS1 condensates act as a selective reservoir for inactive FA proteins. A) Schematic representation of the BioID workflow. U2OS cells with either low or high expression of TNS1-Myc-BioID were treated with biotin, followed by affinity purification of biotinylated proteins and LC-MS/MS analysis. Raw data from four independent experiments were analysed using MaxQuant and statistical significance was assessed using SAINTexpress. Cells with expression of Myc-BioID only were used as a control. B) Gene ontology (GO) cellular component enrichment analysis of proteins identified in TNS1 BioID screen. C) Closeups of GFP-TNS1 condensates in U2OS cells (representative images of the cells in Figure S3F) stained with the indicated antibodies. LC – low contrast; HC – high contrast. Scale bars 5 μm. D) Quantification of relative recruitment of indicated FA components into GFP-TNS1 condensates. The recruitment score is quantified as ratio of integrated density in the condensates and total integrated density (excluding the nuclear signal) normalized on total condensate area in each cell. Quantification was performed from n = 40-68 cells per condition from at least three independent replicates. E) Closeups of GFP-TNS1 condensates in U2OS cells (representative images of the cells in Figure S3G) stained with the indicated phospho-specific antibodies. Scale bars 5 μm. F) Quantification of the relative recruitment of the indicated proteins into GFP-TNS1 condensates as in Figure 2D. Quantification was performed from n = 44-63 cells from at least three independent experiments. Statistical analysis was performed using Kruskal-Wallis test with Dunn’s multiple comparisons test. p < 0.001 (***). G) Representative confocal images of U2OS mGL-TNS1 cells seeded on poly-L-Lysine and stained either with anti-PXN or anti-pPXN antibodies. Nuclei were stained using DAPI. Scale bars 20 μm. H) Expression of GFP-TNS1 in U2OS cells was induced using doxycycline for the indicated times. Representative immunoblots illustrate changes in expression levels of indicated proteins. I) Quantification of relative protein expression from immunoblots in Figure 3H, normalised to protein expression at t = 0 h. Quantification was performed from three independent experiments, data points corresponding to individual replicates are color-coded. Statistical analysis was performed using 2-way ANOVA with Holm-Šídák’s multiple comparison test. Indicated statistical differences are compared to protein expression at t = 0 h timepoint. p < 0.001 (***); ns. – not significant. J) Representative confocal image of U2OS cells expressing GFP-TNS1 stained with anti-DCP1B (mRNA-decapping enzyme 1B) antibody and DAPI. Closeup I. focused on association of TNS1 condensates with P-bodies (white arrowheads), closeup II. highlights the canonical TNS1 localisation in FAs. Profile plot (right) illustrates the relative intensity distribution along the white line indicated in closeup I. Scale bar 20 μm. K) Representative superresolution image (SIM) of U2OS cells expressing GFP-TNS1 stained with anti-EDC3 antibody, phalloidin (F-actin) and DAPI. Maximum intensity projection is shown. Closeup represents a single focal plane. Scale bar 10 μm.

### P-bodies are closely associated to TNS1 condensates

The GO analysis of the BioID screen revealed a significant enrichment of genes related to P-bodies and ribonucleoprotein granules in U2OS cells with high expression of TNS1-Myc-BioID (Figures 3B and S3C). To examine whether TNS1 condensates recruit P-body proteins, we initially stained the GFP-TNS1 expressing U2OS cells for a canonical P-body component^40^ identified in the BioID screen, mRNA-decapping enzyme 1B (DCP1B). Surprisingly, DCP1B staining did not overlap with the GFP-TNS1 condensates but instead localised in their proximity (Figure 3J), suggesting a close association between the P-bodies and TNS1 condensates. We further stained the cells for two well-established P-body markers, enhancer of mRNA-decapping protein 3 (EDC3) and DEAD-Box Helicase 6 (DDX6). Both EDC3 and DDX6 staining recapitulated the close proximity of P-bodies and TNS1 condensates (Figure S3D) which was further confirmed by imaging EDC3 using super-resolved structured illumination microscopy (SR-SIM) (Figure 3K). Since P-bodies are involved in messenger RNA (mRNA) storage, translational repression and mRNA decay, and mRNA recruitment has been reported proximal to force-loaded focal adhesions^41^, we assayed the presence of mRNA in the GFP-TNS1 condensates using fluorescent in situ hybridisation (FISH) with probes targeting mRNA poly-A tails. Although the probes readily detected polyA-mRNA in the cell nucleus and cytoplasm, we did not detect any enrichment of the FISH signal in TNS1 condensates (Figure S3E), indicating that mRNA is either absent from TNS1 condensates or the polyA tails are not accessible for binding of the oligo-dT probe. Overall, these observations further validate the results of our proteomic data, however, the functional significance of P-body association with FA-derived condensates remains the objective of future studies.

### TNS1 condensation is mediated by its intrinsically disordered region

Protein condensation is generally mediated either by site-specific interactions (SSI) between structurally organised domains, by low complexity sequences across intrinsically disordered regions (IDRs), or by a combination of both^37,42^. TNS1 encompasses two N-terminal (protein tyrosine phosphatase, PTP; homologous protein kinase C conserved region 2, C2) and two C-terminal domains (src-homology 2 domain, SH2; phosphotyrosine binding domain, PTB) that are interconnected with a long IDR (Figure 4A). To discriminate between the SSI- and IDR-mediated phase transition mechanisms and to assess how individual regions contribute to TNS1 condensation, we generated a series of deletion mutants lacking each of the individual domains or the entire IDR (Figure 4A). To prevent the potential effect of the endogenous TNS1 on condensation of the mutant variants, we generated a TNS1 knock-out (KO) U2OS cell line. Importantly, TNS1 KO did not alter the expression levels of the associated proteins involved in integrin signalling, such as integrin β1, integrin α5 or PXN (Figure S4A). Expression of the individual GFP-TNS1 mutants in these cells showed that deletion of any of the structurally conserved domains (ΔPTP, ΔC2, ΔSH2 or ΔPTB) did not have any substantial effect on TNS1 condensation. In contrast, deletion of the IDR (ΔIDR) resulted in an almost complete abrogation of the ability of TNS1 to phase separate (Figure 4B). To evaluate whether there are any specific regions within the TNS1 IDR preferentially promoting phase separation, we generated three mutants with sequential deletions of approximately a third of the IDR sequence (ΔIDR1, ΔIDR2 and ΔIDR3) (Figure 4C) and quantified the correlation between relative expression of the individual mutants and their propensity to form condensates. While we observed no significant difference between GFP-TNS1 WT and ΔIDR1, deletion of both IDR2 and IDR3 significantly reduced TNS1 condensation, although to a lesser extent than in the case of ΔIDR (Figures 4D-F). Taken together, we provide evidence that TNS1 phase separation is mediated by its IDR with major contribution from the IDR2 and IDR3 segments.

**Figure 4.**
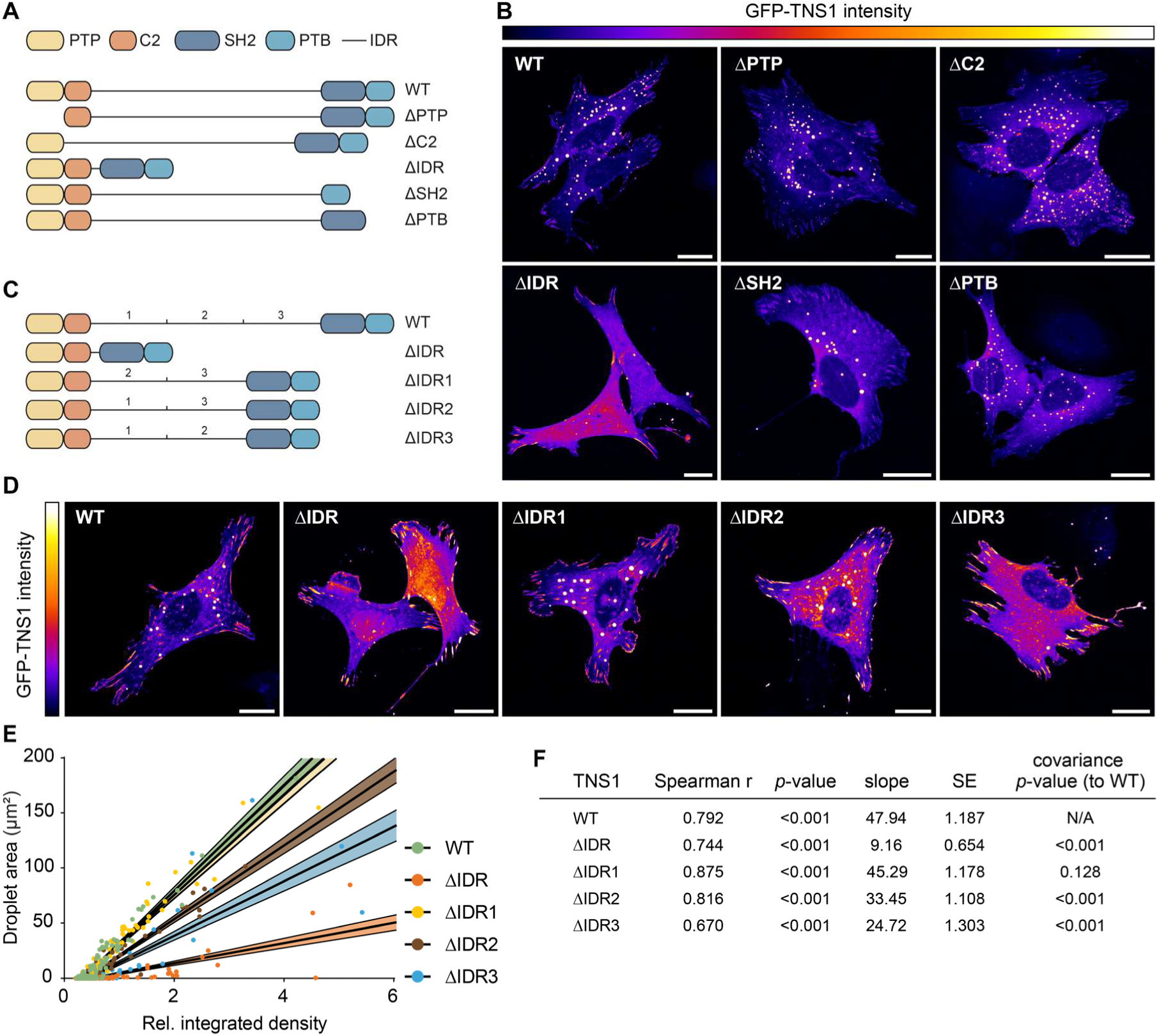
TNS1 condensation is mediated by its intrinsically disordered region. A) Schematic representation of domain organisation of TNS1 WT and its individual deletion mutants. B) Representative confocal images of U2OS TNS1 KO cells expressing indicated GFP-TNS1 variants. Scale bars 20 μm. C) Schematic representation of domain organisation of TNS1 WT and the individual IDR deletion mutants. D) Representative confocal images of U2OS TNS1 KO cells expressing indicated GFP-TNS1 variants. Scale bars 20 μm. E) Correlation between droplet area and relative integrated density quantified from U2OS TNS1 KO cells expressing individual GFP-TNS1 variants. Quantification was performed from n = 90-168 cells per condition. Statistical analysis is shown in Figure S4B. F) Analysis of data from n = 90-168 cells per condition shown in Figure 4E was performed using linear regression analysis, slope and standard error (SE) for each TNS1 variant are shown. Spearman correlation (r) was calculated for each TNS1 variant, Spearman r and the respective p-values are shown. Statistical differences between slopes were assessed by covariance analysis comparing each of the deletion mutants to TNS1 WT.

To further understand the role of the TNS1 IDR in mediating its phase separation, we performed a compositional analysis of the IDR sequence, including its individual segments (IDR1, IDR2 and IDR3) and the respective deletion mutants (ΔIDR1, ΔIDR2 and ΔIDR3). This analysis revealed stronger disorder-promoting hydrophobicity patterns in the sequence corresponding to IDR3, which exhibits the highest disorder propensity (⟨λ⟩) values, as well as higher values of sequence hydrophobicity disorder (SHD) when compared to IDR1 and IDR2 (Figure S4B). Despite its high disorder tendency, IDR3 contains the lowest fraction of charged residues (FCR) among the three IDR subregions and is uniquely characterized by a positive net charge per residue (NCPR), suggesting that the disorder properties of IDR3 derive from compositional factors beyond simple charge repulsion. Furthermore, deletion of IDR3 (ΔIDR3) leads to a marked shift in properties of the remaining IDR sequence, including an increase in aromatic clustering (Ω_aro_) and a fundamental change in sequence charge decoration (SCD) from strongly negative to positive, highlighting the distinctive disorder and compositional properties of this region (Figure S4B). Together, our sequence composition analysis supports our experimental data and suggests that a unique combination of high disorder tendency with specific compositional features of the individual IDR segments play a critical role in modulating the overall phase separation behaviour of TNS1.

### The TNS1 IDR regulates FA dynamics, cell migration and invasion

We next assessed the functional roles of the TNS1 IDR in adhesion dynamics and cell behaviour. To this end, we generated TNS1 KO U2OS cell lines with inducible expression of either GFP alone, GFP-TNS1 WT or GFP-TNS1 ΔIDR, ensuring comparable expression levels between individual cell lines (Figure S5A). First, we assessed the characteristics of TNS1-containing FAs in cells expressing GFP-TNS1 WT or ΔIDR. While we observed no difference in average FA size between TNS1 WT and ΔIDR (Figure S5B), there was a significant reduction in FA area in cells expressing TNS1 ΔIDR (Figure 5A), with a notable, although not statistically significant, increase in FA localisation towards the cell periphery (Figure S5C). In addition, the TNS1 ΔIDR-expressing cells exhibited higher FA assembly and disassembly rates when compared to TNS1 WT cells (Figures 5B and 5C), suggesting a role for the IDR in regulation of FA stability. In line with increased FA dynamics, TNS1 ΔIDR promoted short-lived filopodia tip adhesions and filopodia formation in cell protrusions, localizing into filopodia tips in migrating cells (Figures 5D and S5D). Although compositionally different from FAs, filopodial tip adhesions have been previously shown to probe the cell environment and promote directed cell migration and invasion by maturation into nascent and focal adhesions^43–45^. Similarly, when embedded in 3D fibrillar collagen, cells expressing TNS1 ΔIDR exhibited increased number of short protrusions emanating from the cell body (Figure 5E and S5E), indicating IDR-dependent negative regulatory role of the IDR towards a protrusive phenotype both in 2D and 3D. Further characterisation of cells expressing GFP, GFP-TNS1 WT or GFP-TNS1 ΔIDR, revealed no significant differences in cell area (Figure S5F) or cell shape (Figure S5G), and functional experiments did not show any differences in cell proliferation (Figure S5H), or cell spreading monitored as a change in impedance on FN-coated xCELLigence plates (Figure S5I). Expression of both TNS1 WT and ΔIDR led to a substantial decrease in cell migration when compared to GFP-expressing cells, however, ΔIDR was able to partially revert the inhibitory effect of TNS1 WT on cell migration as evidenced by an increase in both migrated distance (Figures 5F and 5G) and mean cell speed (Figure S5J), with an overall increase in migration directionality (Figure S5K). This correlated with a significantly increased amount of active integrin β1 (Figures 5H and 5I), in line with previously described TNS1 roles in regulating integrin activity^24–27^. In addition, cells expressing either TNS1 WT or ΔIDR exerted higher traction forces on 10kPa polyacrylamide gels than the control cells, but there was no significant difference in traction between the TNS1 variants (Figures 5J and S5L). Finally, expression of both TNS1 WT and ΔIDR led to a significant increase in cell invasion into 3D collagen gels when compared to control cells, with ΔIDR invading substantially more than WT (Figures 5K and 5L). Taken together, we observed a gain-of-function phenotype in TNS1 ΔIDR, suggesting that IDR deletion leads to increased FA dynamics and promotes cell protrusiveness, migration and invasion.

**Figure 5.**
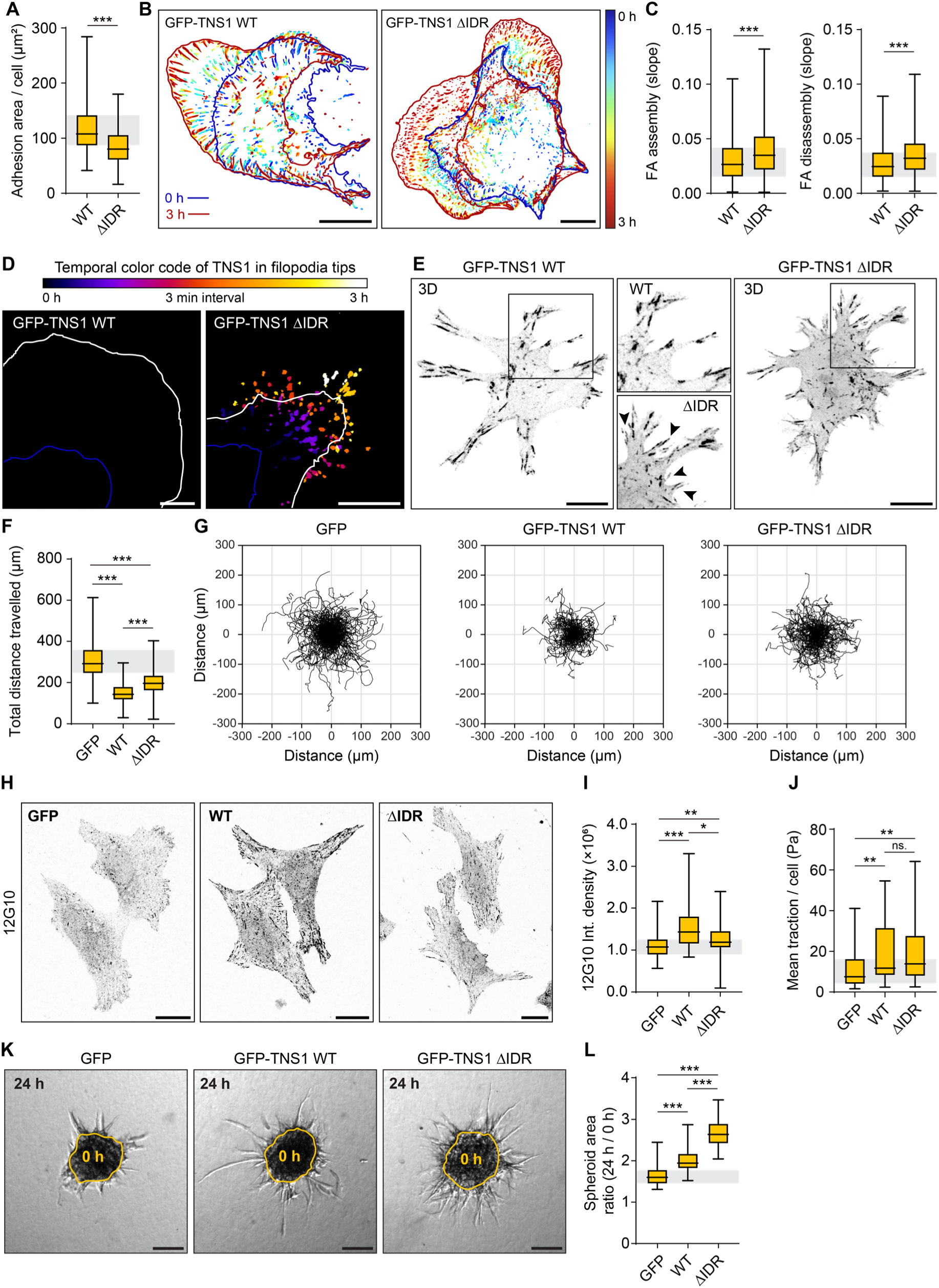
TNS1 IDR regulates FA dynamics, cell migration and invasion. A) Quantification of the FA area of the indicated cell lines from n = 154-165 cells per condition. Statistical analysis was performed using two-tailed Mann-Whitney U test. p < 0.001 (***). B) Representative images of FA dynamics in indicated cell lines. Time-lapse images acquired for 3 h at 3 min intervals were analysed by the Focal adhesion analysis server (FAAS)^93^. Color-coded cell outlines from the initial and the last timepoints are shown. Scale bars 20 μm. C) Quantification of FA assembly (left) and disassembly (right) rates from the indicated cell lines performed using the FAAS^93^. Quantification was performed from n = 41-43 cells per condition analysing n = 3334-4594 (assembly) and n = 3593-5155 (disassembly) individual adhesions. Statistical analysis was performed using two-tailed Mann-Whitney U test. p < 0.001 (***). D) Representative filopodia tip masks from time-lapse images of indicated cell lines migrating on FN-coated coverslips (images shown in Figure S5D). Images were acquired for 3 h at 3 min intervals and filopodia masks from each timeframe have been color-coded as indicated. Outlines of cell protrusions at times t = 0 h (dark blue) and t = 3 h (white) are shown. Scale bars 10 μm. E) Representative 3D projection of cells expressing indicated GFP-TNS1 variants embedded in 3D fibrillar rat tail collagen (the respective multichannel images shown in Figure S5E). Black arrowheads are indicating short TNS1-rich cell protrusions. Scale bars 10 μm. F) Quantification of total distance travelled of n = 159-170 cells per condition over the period of 10 h. Statistical analysis was performed using Kruskal-Wallis test with Dunn’s multiple comparisons test. p < 0.001 (***). G) Cell migration trajectories of cells analysed in Figure 5F. H) Representative images of indicated cell lines stained with 12G10 antibody (active integrin β1). Scale bars 20 μm. I) Quantification of integrated density of the indicated cell lines stained with 12G10 antibody. Quantification was performed from n = 81-94 cells per condition. Statistical analysis was performed using Kruskal-Wallis test with Dunn’s multiple comparisons test. p < 0.05 (*); p < 0.01 (**); p < 0.001 (***). J) Quantification of mean traction force per cell from n = 45-66 cells per condition. Representative images are shown in Figure S5L. Statistical analysis was performed using Kruskal-Wallis test with Dunn’s multiple comparisons test. p < 0.01 (**); ns. – not significant. K) Representative images of 3D spheroid invasion assays using the indicated cell lines. Images at 24 h timepoint are shown with the spheroid outline at t = 0 h shown in yellow. Scale bars 100 μm. L) Quantification of spheroid area ratio at 24 h and 0 h from n = 62-71 spheroids per condition. Statistical analysis was performed using Kruskal-Wallis test with Dunn’s multiple comparisons test. p < 0.001 (***).

### TNS1 condensation is regulated by stress-mediated phosphorylation

Phosphorylation has been repeatedly shown to regulate protein condensation across organisms^16,46–51^. As our screen targeting canonical tyrosine phosphorylation-dependent integrin signalling pathways did not reveal any significant impact on TNS1 condensation, we next focused on Ser/Thr phosphorylation. Since a previous phosphoproteomic study demonstrated that TNS1 undergoes strong Ser/Thr phosphorylation in response to cell stress^52^, we next assessed the potential role of stress-mediated kinase activity in regulating TNS1 condensation using a well-established inducer of oxidative stress, sodium arsenite. Indeed, arsenite treatment of GFP-TNS1-expressing U2OS cells led to a significant translocation of TNS1 from the dense to the dilute phase (cytoplasm) (Figures 6A and 6B, Movies S6-S7), resulting in a substantial decrease in condensate size and area (Figures S6A and S6B). This correlated with a strong arsenite-induced increase in TNS1 Ser/Thr phosphorylation, as well as increased activation of major stress-activated kinases including p38, ERK and AKT (Figures 6C and 6D). To explore this further, we inhibited these kinases with a combination of specific inhibitors (p38- (Doramapimod), MEK- (Trametinib, upstream of ERK) and AKT- (MK2206)) and observed completely impaired GFP-TNS1 translocation to the dilute phase upon arsenite treatment (Figures 6E and 6F, Movie S8), confirming the role of these kinases in TNS1 phosphorylation. As controls, we used U2OS cell lines with inducible expression of either GFP-TNS2, a TNS1 paralog, or GFP fused to Pop-tag, a short protein domain driving cytoplasmic protein condensation^53^. In the case of GFP-TNS2, arsenite stimulation led to a small but significant translocation of the protein from the condensates to the dilute phase, however, its effect was strikingly lower when compared to GFP-TNS1 (Figures S6C, S6D and S6E). In contrast, arsenite treatment of cells expressing GFP-Pop-tag led to a significant increase in condensate formation (Figures S6C, S6D and S6F). Taken together, our results demonstrate that TNS1 condensation is negatively regulated by protein phosphorylation and that this mechanism is protein-specific, rather than a general characteristic of phase separated proteins.

**Figure 6.**
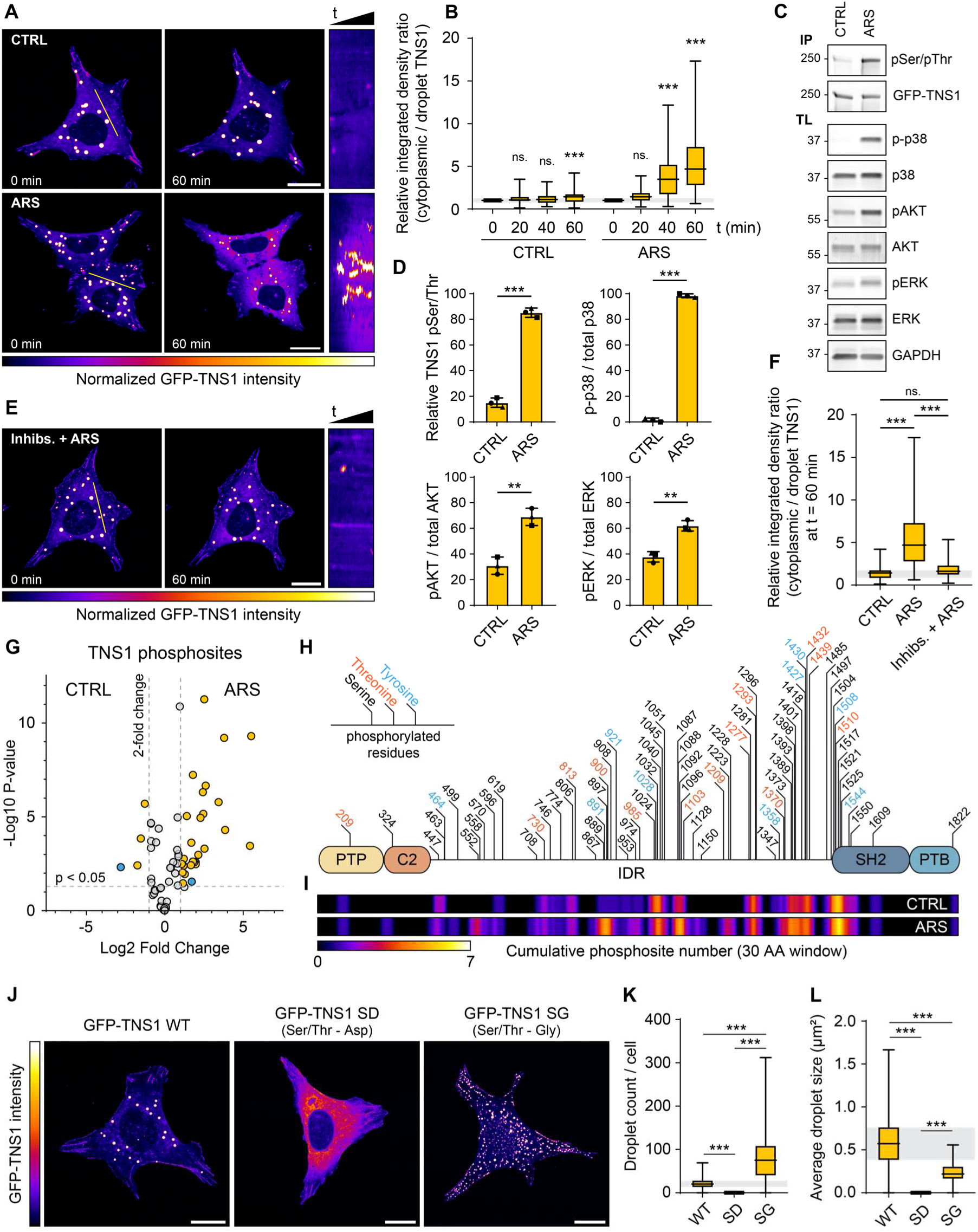
TNS1 condensation is regulated by stress-mediated phosphorylation. A) Representative timelapse images of either non-treated U2OS cells expressing GFP-TNS1 (CTRL) or cells treated with sodium arsenite (ARS). Images were acquired for 60 min at 1 min intervals. Images at t = 0 min and t = 60 min are shown, along with a kymograph of the whole timelapse along the white line. Scale bars 20 μm. B) Quantification of the ratio between the relative cytoplasmic and droplet TNS1 integrated densities at the indicated timepoints from n = 78-115 cells per condition. Statistical analysis was performed using Friedman test with Dunn’s multiple comparisons test comparing the individual timepoints to t = 0 min. p < 0.001 (***); ns. – not significant. C) Representative immunoblots from lysates of either CTRL or ARS-treated cells. GFP-TNS1 was immunoprecipitated to probe with anti-pSer/pThr antibody. D) Quantification of relative TNS1 Ser/Thr phosphorylation and relative activity of p38, ERK and AKT kinases quantified as ratio between phosphorylated and total protein. Quantification was performed from three biological replicates. Statistical analysis was performed using unpaired t-test. p < 0.01 (**); p < 0.001 (***). E) Representative timelapse images of U2OS cells expressing GFP-TNS1 pre-treated with 5 μM Doramapimod (p38), 5 μM Trametinib (MEK1/2) and 5 μM MK2206 (Akt) for 60 min before ARS treatment. Images were acquired for 60 min at 1 min intervals. Images at t = 0 min and t = 60 min are shown, along with a kymograph of the whole timelapse along the white line. Scale bars 20 μm. F) Quantification of the ratio between the relative cytoplasmic and droplet TNS1 integrated densities at t = 60 min from n = 78-115 cells per condition. Statistical analysis was performed using Kruskal-Wallis test with Dunn’s multiple comparisons test. p < 0.001 (***); ns. – not significant. G) Volcano plot of individual TNS1 phosphorylation sites as identified by mass spectrometry of GFP-TNS1 purified from CTRL or ARS-treated cells. Statistically enriched (> 2-fold change at p < 0.05) Ser/Thr phosphosites are indicated in yellow and Tyr phosphosites in blue. H) Schematic representation of localisation of the identified phosphosites within TNS1. I) Representation of the cumulative phosphosite number over 30 amino acid (AA) window within TNS1 purified from CTRL or ARS-treated cells. J) Representative confocal images of U2OS TNS1 KO cell lines with inducible expression of indicated GFP-TNS1 variants. Scale bars 20 μm. K) Quantification of TNS1 droplet count per cell as quantified from n = 339-358 cells per condition. Statistical analysis was performed using Kruskal-Wallis test with Dunn’s multiple comparisons test. p < 0.001 (***). L) Quantification of average TNS1 droplet size per cell as quantified from n = 340-358 cells per condition. Statistical analysis was performed using Kruskal-Wallis test with Dunn’s multiple comparisons test. p < 0.001 (***).

### TNS1 phosphorylation affects IDR charge patterning and prevents condensate assembly

To identify the phosphorylation sites involved in regulating TNS1 phase separation, we next performed a phosphoproteomic analysis of GFP-TNS1 purified from either control (CTRL) or arsenite-treated (ARS) cells (Figures 6G and S6G). In line with our previous results (Figures 6C and 6D), we observed a substantial increase in Ser/Thr phosphorylation in GFP-TNS1 purified from arsenite-treated cells with 26 Ser/Thr sites being significantly enriched (>2-fold) when compared to control (Figure 6G). Interestingly, most of the total of 50 Ser, 13 Thr and 9 Tyr phosphorylated residues identified in our phosphoproteomic data were localized within the TNS1 IDR (Figure 6H), highlighting the importance of this region in TNS1 condensation in response to its phosphorylation state. Moreover, comparing the cumulative phosphosite number over a 30 AA window between control and arsenite-treated conditions revealed several phosphorylation “hotspots” within the IDR (Figure 6I), leading to a dramatic shift in net charge decoration in these regions (Figure S6H) which has previously been shown to affect phase separation in the context of nuclear condensates^54–56^.

We next assessed the condensation propensity of the different TNS1 phosphorylation states. Since our results suggest that TNS1 condensation is regulated by p38-, ERK- and AKT-mediated phosphorylation (Figures 6E and 6F), we analysed the probability of the individual phosphosites to be phosphorylated by these kinases using the Kinase Library tool^57^ (Figure S6I). Based on the prediction, we selected 36 Ser/Thr sites among the constitutively phosphorylated and significantly enriched sites in arsenite-treated cells (Figure S6I, bottom heatmap) and substituted them for either phosphomimicking (Aspartate, TNS1 SD) or unphosphorylatable (Glycine, TNS1 SG) residues. Notably, the substitutions did not have any adverse effect on protein expression, and we were able to generate TNS1 KO U2OS cell lines with inducible expression of individual GFP-TNS1 variants at comparable levels (Figures S6J and S6K). Interestingly, expression of the GFP-TNS1 SD mutant completely impaired the formation of TNS1 condensates. In contrast, the expression of GFP-TNS1 SG led to a substantial increase in TNS1 condensate number and total area per cell, with a decrease in average condensate size when compared to WT (Figures 6J-6L and S6L). Taken together, our results support the role of phosphorylation in the negative regulation of TNS1 condensation in a p38/ERK/AKT-dependent manner.

### TNS1 phosphorylation controls FA dynamics and cell migration

Finally, we characterised the effect of TNS1 phosphorylation on the regulation of FA dynamics and cell behaviour, using the TNS1 KO U2OS cells with inducible expression of either GFP-TNS1 WT, SD or SG as described above. Analysis of FA properties revealed a significant decrease in total FA area in cells expressing TNS1 SD, and a notable, although not statistically significant, increase in TNS1 SG-expressing cells (Figure 7A). In addition, the average FA size was reduced in cells expressing TNS1 SD (Figure 7B) and the FAs were significantly more localised at the cell periphery (Figure 7C), without substantial changes in cell area (Figure S7A) or cell shape (Figure S7B), suggesting a potential role for TNS1 phosphorylation in regulating FA maturation and dynamics. Indeed, analysis of FA dynamics confirmed a significant increase in both FA assembly and disassembly rates in cells expressing TNS1 SD (Figures 7D and 7E), similarly to the cells expressing TNS1 ΔIDR (Figures 5B and 5C). Also, consistent with our observations following TNS1 ΔIDR expression, TNS1 SD cells exhibited an increase in cell migration distance (Figures 7F and 7G) and speed (Figure S7C), with higher cell directionality (Figure S7D) when compared to WT and SG. In contrast, cells expressing TNS1 SG showed a marked reduction in cell migration distance (Figures 7F and 7G) and speed (Figure S7C). Together, our data suggest a role of TNS1 phosphorylation in regulating FA dynamics and maturation, with direct implications for adhesion dynamics and cell migration.

**Figure 7.**
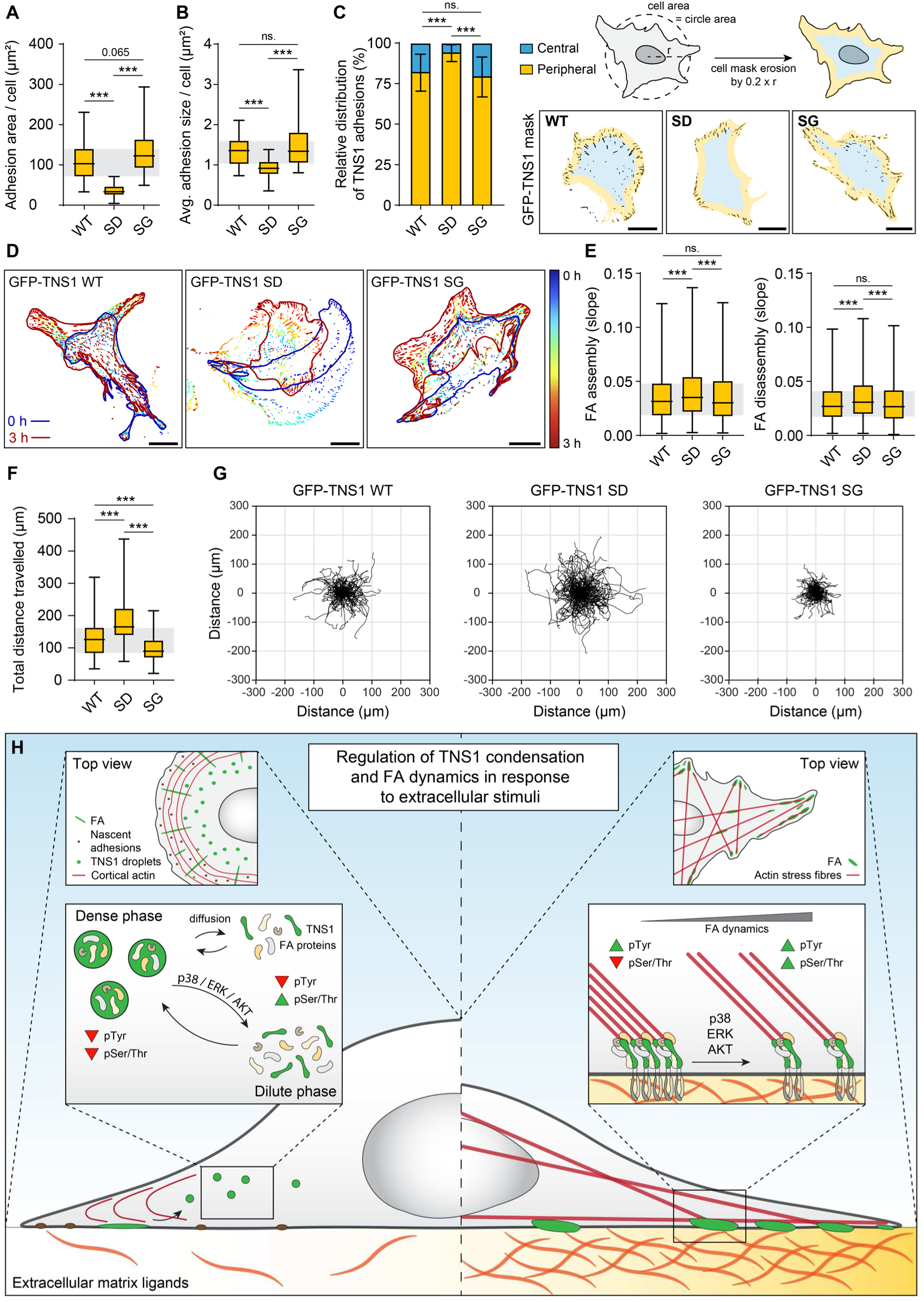
TNS1 phosphorylation controls FA dynamics. A) Quantification of adhesion area per cell from n = 31-42 cells per condition. Statistical analysis was performed using one-way ANOVA with Holm-Šídák’s multiple comparison test. p < 0.001 (***). B) Quantification of average adhesion size per cell from n = 32-42 cells per condition. Statistical analysis was performed using Kruskal-Wallis test with Dunn’s multiple comparisons test. p < 0.001 (***); ns. – not significant. C) Quantification of relative distribution of peripheral and central FAs from the indicated cell lines. Analysis was performed from n = 30-42 cells per condition by eroding the cell mask by the factor of 0.2 × radius (r) of a circle with area corresponding to the area of the cell. Representative images corresponding to central and peripheral areas with GFP-TNS1 adhesion masks are shown. Scale bars 20 μm. Statistical analysis was performed using 2-way ANOVA with Holm-Šídák’s multiple comparison test. p < 0.001 (***); ns. – not significant. D) Representative images of FA dynamics in indicated cell lines. Time-lapse images acquired for 3 h at 3 min intervals were analysed by the Focal adhesion analysis server (FAAS)^93^. Color-coded cell outlines from the initial and the last timepoints are shown. Scale bars 20 μm. E) Quantification of FA assembly (left) and disassembly (right) rates performed using the FAAS^93^. Quantification was performed from n = 30-42 cells per condition analysing n = 1900-1958 (assembly) and n = 1439-2234 (disassembly) individual adhesions. Statistical analysis was performed using Kruskal-Wallis test with Dunn’s multiple comparisons test. p < 0.001 (***); ns. – not significant. F) Quantification of total distance travelled of n = 134-153 cells per condition over the period of 10 h. Statistical analysis was performed using Kruskal-Wallis test with Dunn’s multiple comparisons test. p < 0.001 (***). G) Cell migration trajectories of cells analysed in Figure 7F. H) Model of regulation of TNS1 condensation and FA dynamics in response to extracellular cues. Limited availability of integrin ECM ligands promotes formation of TNS1 condensates that selectively buffer the local concentration of inactive FA components. TNS1 condensates dynamically exchange material with the surrounding cytoplasm by passive diffusion or disassemble upon TNS1 phosphorylation by stress-activated kinases. In contrast, on ECM-rich substrates, stable adhesions are formed. Phosphorylation of TNS1 in these leads to increased FA dynamics promoting directed cell migration.

## Discussion

In this study, we characterized the role of TNS1 condensation in regulating FA dynamics. Through comprehensive validation, we demonstrated that TNS1 undergoes concentration-dependent phase separation in cells, including at endogenous protein levels. Using proximity biotinylation proteomics, we identified the principal components of TNS1 condensates and validated their role in maintaining inactive FA protein pools. Furthermore, we identified the TNS1 IDR as a region driving TNS1 condensation and established its role in regulating TNS1-mediated biological functions. Finally, we demonstrated that TNS1 phosphorylation by stress-activated kinases negatively regulates TNS1 condensate formation, as well as FA dynamics and cell migration (Figure 7H). Our data indicate that stress-signals can act as physiologically relevant triggers for release of adhesion proteins from cytoplasmic reservoirs to rapidly support new adhesion formation.

The complexity of FA organisation and their dynamic regulation has prompted views that FAs should be characterized as phase separated molecular assemblies^5,6^. Indeed, a number of individual FA components, including GIT, PIX^18^, FAK^14^, CAS^14,58^, LIMD1^16^, PXN^19^, TLN^5^, KANK1^17^ and TNS1^15^, have been observed to form condensates *in vitro* and/or in cells, either alone or in combination with other FA components. Intriguingly, we demonstrated that TNS1 condensates are selective in nature, recruiting a subset of FA components rather than all components non-specifically. Considering the previous findings, our results suggest a potential hierarchical organisation within FA-derived condensates, however, systematic studies evaluating protein recruitment to individual condensates are required to address this aspect comprehensively.

We demonstrated that a local increase in endogenous TNS1 concentration upon FA disassembly or lack of integrin-ECM engagement in both 2D and 3D conditions results in the formation of TNS1 condensates. This is in line with the proposed functions of phase separation in buffering local biomolecule concentrations into compartmentalised systems^8,37,46,59^. The sequestering role of TNS1 condensates is also supported by our results suggesting that TNS1 phase separation can acutely increase the amounts of recruited proteins, possibly by protecting them from degradation. Together, our observations address a longstanding question of how expression levels and local availability of FA components are maintained in the proximity of adhesion sites.

We provided evidence that TNS1 condensation is driven by its IDR. This is particularly interesting in the context of differences between the individual tensin paralogs. While the structurally ordered domains at the N- and C-terminus of TNS1-3 are highly conserved, their IDRs share very little sequence homology^20^, implying their importance in regulating some of the distinct functions of the tensin family members^20,21^. In the case of TNS1, the IDR seems to play an inhibitory role towards the promigratory and protrusive phenotype stimulated by the conserved domains. It remains a focus of future studies to determine whether the same regulation holds true also for the other tensin family members.

We further demonstrate that predominantly IDR-targeted TNS1 phosphorylation by stress-activated kinases p38, ERK and AKT, regulates TNS1 condensation and FA dynamics. In contrast to Tyr phosphorylation largely associated with active integrin signalling, Ser/Thr phosphorylation of FA proteins is far less understood. Although tightly balanced levels of CDK1-mediated phosphorylation are essential for maintaining FA stability^60–62^, hyperphosphorylation has been widely linked to FA disassembly, e.g. by disrupting FAK/p130Cas interactions within FA complexes^63,64^ or by targeting phosphorylated kindlin for degradation before mitotic entry^65^. Interestingly, although TNS1 harbours 312 Ser/Thr residues, phosphomimetic substitution of only 36 of them had a striking effect on TNS1 condensation and its FA-mediated biological functions. Moreover, since p38/ERK MAPKs and CDK1 kinases exhibit a strong preference for Ser/Thr-Proline motifs, it is plausible to hypothesize that they share at least some of the phosphorylation sites within TNS1, providing an additional level of regulatory control in key biologically relevant processes, such as the essential steps of adhesion disassembly and assembly during cell division^66,67^.

It is important to note that the stress-responsive kinases are activated by diverse physiological stimuli that, among other processes, drive EMT and cancer progression. For instance, TGFβ acts as a potent activator of both p38 and ERK MAPKs, as well as AKT^68,69^. In line with the role of TGFβ in promoting EMT and cell migration^70^, expression of the TNS1 mutant mimicking phosphorylation by these kinases resulted in increased FA dynamics and migratory cell behaviour consistent with more mesenchymal phenotype. Therefore, phosphorylation-controlled TNS1 condensates acting as FA protein reservoirs may represent a previously unknown level of control of cell morphology changes linked to EMT. Beyond mesenchymal cells and cancer, TNS1 is enriched in neutrophils and monocytes (Human Protein Atlas). While its role in these cells remains largely unexplored, our data may explain early observations of TNF-mediated p38 activation acting as a stop signal that promotes firm neutrophil adhesion and inhibits chemotaxis towards interleukin-8^71^. Furthermore, kinase-dependent regulation of TNS1 could contribute to the crosstalk between integrin and MAPK signalling that has been demonstrated in multiple physiological contexts. Acute integrin β1-dependent p38 activation was observed in human osteosarcoma cells^72^ and in cardiomyocytes subjected to mechanical stress^73,74^. In addition, interleukin-8 production in natural killer cells has been shown to be dependent on integrin β1-mediated p38 activation^75^. Our results therefore support existence of a regulatory feedback loop in which acute integrin-mediated activation of stress-associated kinases leads to phosphorylation of FA components, thereby increasing FA dynamics and promoting cell migration.

## Limitations of the study

During the implementation of this study, we encountered certain technical limitations. Although we aimed to complement our validation of TNS1 condensation with *in vitro* data using purified TNS1, due to the large size of the protein and degradation issues during purification, we were unable to obtain sufficient material from insect or mammalian expression systems. Nevertheless, we provided compelling evidence supporting TNS1 condensation within living cells. Finally, some of our interesting findings were beyond the scope of this study. For instance, we validated that TNS1 condensates closely associate with P-bodies, in line with previous (high-throughput) FA interactome screens that identified some of the P-body components^58,76,77^. In addition, we detected multiple centrosomal proteins in our BioID screen. This is not surprising, since centrosomes act as biomolecular condensates^78^, and other FA proteins, such as paxillin, have been previously shown to localise to centrosomes by us^79^ and others^80^. However, both the functional significance of P-body association with FA-derived condensates and the role of FA proteins in centrosomal assembly and cell polarity remain the focus of future studies.

## Materials and Methods

### Plasmids and cloning

The plasmid encoding pEGFP-TNS1 WT has been described previously^28^. For simplicity, abbreviation GFP instead of EGFP is used throughout the study, except in the plasmid names. The TNS1 isoform used in this study corresponds to the canonical UniProt entry Q9HBL0-3 missing the amino acids 1-125. In case of phosphorylation sites, we always use the numbering corresponding to full Q9HBL0-3 isoform as annotated on phosphosite.org.

Individual TNS1 mutants (ΔPTP, ΔC2, ΔIDR, ΔSH2, ΔPTB, ΔIDR1, ΔIDR2, ΔIDR3) were generated by a whole plasmid synthesis approach using homemade *Pfu-X7* high fidelity polymerase^81^, pEGFP-TNS1 WT as a template and the respective primers (listed in Table S1). PCR was carried out for 18 cycles and the reactions were purified using KAPA Pure Beads (Roche), digested with *DpnI* (Thermo Fisher Scientific) and transformed into *E. coli* DH5α competent bacteria. pEGFP-TNS1 SD and SG were generated by NEBuilder® HiFi DNA Assembly (New England Biolabs) of the commercially synthesized coding sequences (TNS1 SD_1/2, TNS1 SG_1/2, sequences listed in Table S1, GenScript) and the backbone of pEGFP-TNS1 WT cleaved by *XhoI*/*SalI*. All the constructs were verified by analytical digestion and sequencing.

pSB EGFP-TNS1 WT/ΔIDR/SD/SG constructs were generated by cloning from pEGFP-TNS1 WT/ΔIDR/SD/SG into the *SleepingBeauty* transposon vector (pSB MCS) via *AgeI*/*XhoI* and *AgeI*/*SalI* sites, respectively.

TNS1 BioID construct was generated by NEBuilder® HiFi DNA Assembly (New England Biolabs). TNS1 WT cDNA was PCR-amplified from pEGFP-TNS1 WT using *Pfu-X7* polymerase^81^ with the respective primers (TNS1 Hifi F/R) and assembled into pPB-BirA* vector^82^ cleaved with *BspEI*.

### Cell lines, cell culture and transfection

Human osteosarcoma cell line, U2OS, has been used throughout the study. The cells were cultured in Dulbecco’s Modified Eagle’s Medium supplemented with 10% fetal bovine serum (FBS), 100 U/ml Penicillin, 0.1 mg/ml Streptomycin and 2 mM L-Glutamine. The cells were kept in a humidified incubator with 5% CO_2_ at 37 °C. Cell transfection was performed using Lipofectamine 3000 (Invitrogen), siRNA was transfected using Lipofectamine RNAiMAX (Invitrogen) following the manufacturer’s instructions.

U2OS cell lines with inducible expression of individual GFP-TNS1 variants were generated by co-transfecting the cells with pSB EGFP-TNS1 (WT/ΔIDR/SD/SG) plasmid together with a vector encoding hyperactive SB transposase (pSB100X)^83^ in a 5:1 ratio. Ten days after transfection, expression of GFP-TNS1 was induced by adding 0.25 μg/ml doxycycline for 48 h and the cells were sorted using SONY SH800S Cell Sorter for GFP fluorescence. After expansion, the cells were sorted one more time to achieve cell population with uniform expression. Before each experiment, unless indicated, GFP-TNS1 expression was induced for at least 48 h by 0.25 μg/ml doxycycline.

U2OS cell lines stably expressing TNS1-BioID were generated by co-transfecting the cells with pPB TagBFP-T2A-TNS1-Myc-BioID plasmid together with a vector encoding a hyperactive transposase (pCMV-hyPBase)^84^ in a 4:1 ratio. Ten days after transfection, the cells were sorted based on TagBFP fluorescent signal into two populations with either low or high expression. This correlated with TNS1-Myc-BioID expression as validated by IF and western blot.

### Generation of TNS1 knock-out cell line

The CRISPR/Cas9 guide sequence (TGAGAGCGGAACATACCGCT) targeting Exon 17 of TNS1-201 transcript (Transcript ID - ENST00000171887.8) was designed using CRISPOR ^85^ and cloned into pX330 U6-Chimeric hSpCas9-Venus plasmid ^86^ via *BbsI* (New England Biolabs) sites. U2OS cells were transfected with the corresponding plasmid and, 48 h after transfection, Venus-positive cells were sorted using SONY SH800S Cell Sorter. Single cell clones were raised from the sorted population and knock-out efficiency was determined using immunoblot with anti-TNS1 antibody. For further validation, genomic DNA from 5 TNS1 knock-out (KO) clones and 5 clones with unaffected TNS1 expression levels was isolated and used as a template to PCR-amplify a 566 bp-long fragment surrounding the cleavage site (F: GGGACACTCTGGGACTTGTG; R: GTCCTACCCCATGGAGCCTA) that was sequenced. Validated clones were pooled into TNS1 KO and TNS1 ctrl populations.

### TNS1 endogenous tagging

U2OS cells with endogenously tagged TNS1 were generated using homology-directed repair-mediated (HDR) CRISPR/Cas9 targeting Exon 6 of TNS1-201 transcript (Transcript ID - ENST00000171887.8). The corresponding guide sequence (CATGAGTGTGAGCCGGACCA) was cloned into pXPR_001 lentiCRISPR v1 plasmid (Addgene #49535) cleaved with *BsmBI* (NEB). Final construct (pXPR_001-sgTNS1) was verified by sequencing. Template for HDR was designed based on a double-cut dsDNA donor approach^87^. The sequence of a bright monomeric fluorescent protein, mGreenLantern^38^, followed by a short linker in frame with the TNS1 open reading frame was surrounded by ca. 850 base pairs-long homology arms. The whole cassette was flanked with sequences recognized by the TNS1 Cas9 guide to generate double-cut dsDNA donor for the HDR. The sequence was synthesized into pUC57mini cloning vector (pUC57-TNS1 HDR, sequence listed in Table S1, GenScript). U2OS cells were co-transfected with pXPR_001-sgTNS1 and pUC57-TNS1 HDR plasmids in 1:4 ratio using Lipofectamine 3000 (Invitrogen). 5 days after transfection, the cells were sorted using SONY SH800S Cell Sorter for green fluorescence and re-sorted again after expansion to obtain cell population with the highest fluorescent signal. The specificity of TNS1 endogenous tagging was verified using IF with a TNS1-specific antibody and siRNA knock-down of TNS1 (Figures S2B-D).

### Image acquisition

Fixed samples were imaged using Marianas spinning disk confocal microscope with a Yokogawa CSU-W1 scanning unit and equipped with an Orca Flash4 sCMOS camera (Hamamatsu Photonics). The objectives used were 40×/1.1 NA W LD C-Apochromat (Zeiss) and 63×/1.4 NA O PLAN-Apochromat (Zeiss). Images were acquired using SlideBook 6 software (Intelligent Imaging Innovations). For imaging of live cells, the samples were maintained at 37 °C and 5% CO_2_ in a stage top humidified incubator (Okolab). Cells embedded in 3D collagen were imaged using Leica Stellaris 8 Falcon FLIM microscope with HC PL APO 86×/1,20 NA W motCORR objective. Images were acquired using LAS X software. 3D invasion assays were imaged using Leica Thunder equipped with 5×/0.12 NA N PLAN objective and Leica K8 camera. Images were acquired using LAS X software with Navigator. Cell migration assays were imaged on Nikon Eclipse Ti2-E equipped with 10×/0.3 NA Nikon CFI PLAN-Fluor objective and Hamamatsu sCMOS Orca Flash 4.0 camera. Images were acquired using NIS-Elements AR 5.11.01 software. The samples were maintained at 37 °C and 5% CO_2_ in a stage top humidified incubator (Okolab).

Super-resolved 3D structural illumination microscopy (SR-SIM) was performed using Deltavision OMX V4 (Applied Precision, GE Healthcare) equipped with a 60×/1.42 NA SIM PLAN Apo N (Olympus) objective and 3x PCO edge Front illuminated sCMOS camera. Images were acquired using OMX Acquistion v3.70 software and SIM reconstruction was performed in softWoRx Deconvolution v7.0.0. Chromatic shift of the reconstructed SIM images was corrected using Chromagnon v0.91^88,89^ based on 0.2 μm TetraSpeck microsphere (Invitrogen) data.

### Immunofluorescence

Cells were seeded on ECM-coated glass coverslips either for 3-4 h or overnight as indicated and fixed for 10 min in pre-warmed 3% PFA (Thermo Fisher Scientific) in PBS at 37 °C. Fixed cells were washed with PBS three times for 5 min at RT. When needed, cells were permeabilised using a permeabilization buffer containing 10% horse serum (Gibco) and 0.3% Triton X-100 Surfact-Amps (Thermo Fisher Scientific) in PBS for 10 min at RT, followed by three 5 min washes with PBS. Primary and secondary antibodies were incubated in 4% horse serum (Gibco) in PBS for 1 h at RT, with three washes with PBS between and after the respective incubations. The antibodies used for IF were as follows: anti-TNS1 (Sigma, SAB4200283); anti-FAK (BD Transduction Laboratories, 610088); anti-p130CAS (Cell Signalling Technologies (CST), 13846S); anti-ZYX (Abcam, ab109316); anti-KANK2 (Sigma, HPA015643); anti-TLN1 (Novus Biologicals, NBP2-50320); anti-TLN2 (Novus Biologicals, NBP2-50322); anti-VCL (Sigma, V9131); anti-PXN (GeneTex, GTX125891); anti-DCP1B (CST, 13233S); anti-DDX6 (Bethyl Laboratories, A300-460A); anti-EDC3 (Abcam, ab168851); anti-pY (BD Transduction Laboratories, 610000); anti-pCAS Y410 (CST, 4011S); anti-pFAK Y397 (CST, 8556S); anti-pPXN Y118 (CST, 2541S); anti-active ITGB1 clone 12G10 (isolated from hybridoma cells, AB_928074); anti-Myc-Tag (CST, 2276S); anti-TNS3 (Rb33; gift from K. Clark, University of Leicester, Leicester, England, UK^21^). Streptavidin conjugated with AlexaFluor680 was used to detect protein biotinylation. DAPI (Life Technologies) was used to detect cell nuclei, and either Phalloidin Atto647N (Sigma) or Spy-FastAct (Spirochrome) probes were used to stain Actin. WGA AlexaFluor647 was used to stain cell membranes. Alexa Fluor–conjugated secondary antibodies (Alexa Fluor 488-, 555-, 568- and 647-conjugated anti-mouse and anti-rabbit; Thermo Fisher Scientific) were used.

### Western blotting

Protein extracts were resolved using SDS-PAGE (4-20% gradient, BioRad) under denaturing conditions and protein were transferred to low fluorescence Immobilon®-FL PVDF membrane (Milipore). Membranes were blocked using AdvanBlock-Fluor blocking solution diluted with PBS in 1:1 ratio. Antibodies were diluted in the blocking solution at indicated concentrations, primary antibodies were incubated overnight at 4 °C, secondary antibodies were incubated for 1 h at RT. Primary antibodies used were as follows: anti-GFP (1:5000, Abcam, ab290); GAPDH (1:4000, HyTest, 5G4 MAb 6C5); anti-TNS1 (1:1000, Sigma, SAB4200283); anti-ZYX (1:1000, Abcam, ab109316); anti-PXN (1:1000, GeneTex, GTX125891); anti-TLN1 (1:1000, Novus Biologicals, NBP2-50320); anti-VCL (1:1000, Sigma, V9131); anti-Myc-Tag (1:1000, CST, 2276S); anti-ITGB1 (1:1000, BD Transduction Laboratories, 610468); anti-ITGA5 (1:1000, Invitrogen, PA5-82027); anti-pSer/Thr (1:1000, BD Transduction Laboratories, 612549); anti-p38 (1:1000, CST, 9212S); anti-p-p38 (1:1000, CST, 9216S); anti-AKT (1:1000, CST, 2920S); anti-p-AKT (1:1000, CST, 9275S); anti-ERK (1:1000, CST, 4696S); anti-p-ERK (1:1000, CST, 4370S). Streptavidin AlexaFluor680 was used to detect protein biotinylation. Relevant AzureSpectra Fluorescent Secondary antibodies with 490, 650 and 800 labels were used at 1:3000 dilution throughout the study. Membranes were washed using TBST in between and after antibody incubations and scanned using Sapphire FL Biomolecular Imager (Azure Biosystems).

### Compound screen

U2OS mGL-TNS1 cells were seeded at FN-coated glass-bottom 24-well plates, left to adhere overnight and treated with either DMSO or indicated compounds as follows: Blebbistatin (25 μM, 60 min, StemCell Technologies, 72402); Calpeptin (50 μM, 30 min, Selleckchem, S7396); H1152 (5 μM, 120 min, MedChemExpress, HY-15720); Lapatinib (0.5 μM, 120 min, MedChemExpress, HY-50898); LY294002 (10 μM, 60 min, Selleckchem, S1105); MG132 (10 μM, 4 h, MedChemExpress, HY-13259); Nocodazole (10 μM, 4 h, MedChemExpress, HY-13520); Ro3306 (20 μM, 60 min, MedChemExpress, HY-12529); Saracatinib (1 μM, 60 min, Selleckchem, S1006); Vanadate (1 mM, 30 min, Sigma, 5086050004); Y27632 (10 μM, 60 min, MedChemExpress, HY-10071). The cells were fixed, and stained using WGA and DAPI to detect cell membrane and nuclei, respectively.

### Fluorescence recovery after photobleaching (FRAP)

U2OS cells expressing GFP-TNS1 were imaged using 3i spinning disk microscope with 1 s interval between timeframes. A z-stack of 9 images (0.27 μm step) was acquired for 9 s before and for 100 s after photobleaching one GFP-TNS1 droplet per cell. Bleaching was performed with 1 ms laser burst across all z-stacks. The acquired timelapse images were analysed in Fiji (ImageJ). First, the maximum intensity projection of the whole field of view was corrected for bleaching (histogram matching) and the individual photobleached droplets were then analysed separately in a cropped ROI. The individual timepoints were aligned using the droplet as a landmark to account for droplet movement during analysis. The intensity spectra of photobleached droplets were combined, normalized to min-max values and a batch FRAP fit was applied to all spectra.

### Correlative light-electron microscopy (CLEM)

U2OS pSB GFP-TNS1 cells induced with doxycycline for 48 h were seeded on fibronectin-coated gridded glass bottom dish (MatTek) and left to adhere overnight. Cells were fixed with warm 4% PFA (Thermo Fisher Scientific) in 0.2 M HEPES (pH 7.4) for 10 min at 37°C and washed twice with 0.2 M HEPES. Cells were stained with DAPI and a confocal z-stack was acquired using 3i CSU-W1 spinning disk microscope with 63× objective as described above. Subsequently, the cells were fixed with 2% glutaraldehyde (Sigma) in 0.2 M HEPES for 2 h at RT and washed twice with 0.2 M HEPES. The samples were than post-fixed with 1% osmium tetroxide and 1.5% potassium ferrocyanide solution, dehydrated in Ethanol and embedded in Epon resin. Thin sections (70 nm) were cut on an ultramicrotome, collected to single-slot electron microscopy copper grids, and stained with 1% uranyl acetate and 0.3% lead citrate solutions. The samples were imaged at 80 kV on Jeol JEM-1400 Plus transmission electron microscope. 19 electron micrographs of the cell of interest at 4000× magnification were stitched together using TrakEM2 plugin in Fiji (ImageJ). The resulting montage was overlaid with maximum intensity projection of 3 bottom slices (z-step 0.27 μm) of the confocal stack using the ec-CLEM plugin in Icy (https://icy.bioimageanalysis.org/). The images were correlated with non-rigid transformation using GFP-TNS1 condensates as landmarks. High magnification images of GFP-TNS1 condensates were imaged at 20000×.

### Immunoprecipitation

U2OS cells expressing GFP-tagged proteins were seeded on one 10 cm plate per condition (15 cm for MS). Cells were washed twice with ice-cold PBS, scraped into residual PBS, transferred into 1.5 ml tube, and centrifuged at 400×g for 2 min at 4°C. Cell pellet was resuspended in 500 μl of lysis buffer containing 75 mM Tris (pH 7.4), 200 mM NaCl, 2 mM MgCl_2_, 1.5% Triton X-100 and 0.75% NP-40, supplemented with Complete protease inhibitors mix (Roche), PhosStop phosphatase inhibitors mix (Roche), and GENIUS Nuclease (Santa Cruz) in recommended concentrations. The lysates were incubated on ice for 20 min with occasional gentle vortexing to facilitate cell lysis and centrifuged at 13000×g for 10 min at 4°C. Part of the cleared lysates was heated with 8× reducing sample buffer to generate total lysate samples. The remining lysate was incubated for 2 h at 4°C with GFP-Trap Magnetic Agarose (Chromotek) previously washed twice with a wash buffer containing 50 mM Tris pH 7.4, 150 mM NaCl and 1% NP-40. After the incubation, the beads were washed once with lysis buffer, twice with wash buffer, eluted into 2× reducing sample buffer and analysed by SDS-PAGE and immunoblot. Samples for MS analysis were washed additional three times with TBS (50 mM Tris pH 7.4, 150 mM NaCl) and stored dry at −80°C until further processed.

### BioID sample preparation

The sample preparation for BioID was carried out as described previously ^90^. The U2OS cell lines stably expressing Myc-BioID only, and with low and high expression of TNS1-Myc-BioID were seeded on two 10 cm plates and supplemented with biotin to a final concentration of 50 μM for at least 18 h. After labelling, the cells were washed twice with PBS and harvested into lysis buffer containing 50 mM Tris (pH 7.4), 500 mM NaCl, 0.2% SDS, 1 mM DTT and protease inhibitors (Roche). Lysates were transferred to a tube containing 20% TritonX-100 (final concentration 2%), passed four times through a 21G needle and 50 mM Tris pH 7.4 (360 μl per plate) was added to the tubes. Subsequently, the lysates were passed additional four times through a 27G needle and centrifuged at 16000×g for 10 min at 4°C. Biotinylated proteins were captured from the cleared lysates by overnight rotation with magnetic Streptavidin beads (ReSyn Biosciences). After incubation, the beads were washed (5 min at RT per wash) twice with 2% SDS, once with a buffer containing 0.1% sodium deoxycholate, 1% Triton X-100, 500 mM NaCl, 1 mM EDTA and 50 mM HEPES (pH 7.5), and once with buffer containing 0.5% NP-40, 0.5% sodium deoxycholate, 1 mM EDTA and 10 mM Tris.Cl (pH 7.4). Proteins were eluted in 2× reducing sample buffer saturated with biotin, heated at 70°C for 10 min and subjected to SDS-PAGE, followed by reduction, alkylation and in-gel digestion. Samples from four independent biological replicates were analysed by MS.

### Mass spectrometry and data analysis

The LC-ESI-MS/MS analyses were performed on a nanoflow HPLC system (Easy-nLC1200, Thermo Fisher Scientific) coupled to the Exploris 480 mass spectrometer (Thermo Fisher Scientific) equipped with a nano-electrospray ionization source and FAIMS interface. Digested peptide samples were dissolved in 0.1% formic acid and injected for analysis. Wash runs were submitted between samples to reduce peptide carry-over. Peptides were first loaded on a trapping column and subsequently separated inline on a 15 cm C18 column (75 μm x 15 cm, ReproSil-Pur 3 μm 120 Å C18-AQ, Dr. Maisch HPLC GmbH). The mobile phase consisted of water with 0.1% formic acid (solvent A) and acetonitrile/water (80:20 (v/v)) with 0.1% formic acid (solvent B).

For BioID samples, a 60 min non-linear gradient was used to elute peptides (50 min from 5% to 21% solvent B followed by 22 min from 21 % to 36 min solvent B and in 5 min from 36% to 100% of solvent B, followed by 5 min wash stage with solvent B). MS data were acquired automatically using Thermo Xcalibur 4.1 software (Thermo Fisher Scientific). A data dependent acquisition method consisted of repeated cycles of one MS1 scan covering a range of m/z 300 – 1750 plus a series of HCD fragment ion scans (MS2 scans) for the most intense peptide ions from the MS1 scan. Raw data files were analysed using MaxQuant software (version 2.0.3.0) and the MS/MS spectra were searched against a human database (Uniprot-SwissProt, version 230427) using Andromeda search engine. Search parameters were set to trypsin digestion with two missed cleavages, carbamidomethyl (C) as a fixed modification, and oxidation (M), acetyl (protein N-term), biotinylation (K) and biotinylation (protein N-term) as variable modifications. The identified peptides were filtered based on FDR 0.01. Unique peptide counts identified for each protein group were subjected to statistical analysis using SAINTexpress (Significance Analysis of INTeractome) ^39^ in SPC mode with default settings (n-iter 4000, lowMode off, minFold on, normalize on) and the results were visualized using ProHits-viz ^91^. Raw MS data are available through PRIDE repository^92^ (to be specified) and through Zenodo (https://zenodo.org/communities/tensinaction). Processed data are available as a supplementary file (Table S2).

For GFP-TNS1 IP samples, a 120 min gradient was used (62 min from 5 % to 21 % solvent B followed by 48 min from 21 % to 36 % solvent B and in 5 min from 36% to 100% of solvent B, followed by 5 min wash stage with solvent B). Samples were analysed by a data independent acquisition (DIA) LC-MS/MS method. MS data were acquired automatically using Thermo Xcalibur 4.6 software (Thermo Fisher Scientific). In the DIA method, a duty cycle contained a full scan (m/z 395 – m/z 1005) and 30 DIA MS/MS scans covering the m/z range of 400-1000 with variable width isolation windows. Raw data from 4 independent replicates were analysed in Spectronaut (Biognosys, version 19.2.). DirectDIA approach was used to identify proteins against human protein database (Uniprot-SwissProt, version 2409) and label-free quantifications were performed using MaxLFQ. Fixed modifications were set to carbamidomethyl (C), variable modifications were set to oxidation (M), acetyl (protein N-term) and phospho (STY). The identified peptides were filtered based on FDR 0.01 and phosphopeptides were filtered based on localisation probability set to 0.7. Normalisation on GFP-TNS1 quantity was performed and the differential abundance of the identified phosphosites was tested using unpaired t-test. Raw MS data are available through PRIDE repository^92^ (to be specified) and through Zenodo (https://zenodo.org/communities/tensinaction). Processed data are available as a supplementary file (Table S3).

### Functional assays

#### Proliferation

The indicated cell lines were seeded in a technical triplicate at 8,000 cells/96 well plate well and allowed to grow for 48 hours. Proliferation was assessed using the AlamarBlue HS reagent (Thermo Fisher Scientific) according to the manufacturer’s instructions. Three independent biological replicates were performed.

#### Cell spreading

Cell spreading was assessed using xCELLigence RTCA eSight (Agilent). 15,000 cells per well were seeded in technical triplicates into a fibronectin-coated E-Plate View 96 (Agilent). Changes in impedance were monitored for 1 h after seeding the cells at 10 min intervals and normalised to the first timepoint. Three independent biological replicates were performed.

#### Focal adhesion dynamics

Cells were seeded on fibronectin-coated µ-Slide 8 Well (Ibidi) and allowed to adhere for 3 h. Individual cells were imaged for 3 h at 3 min intervals, acquiring a z-stack of 5 stacks with 0.5 μm spacing. During image acquisition, the cells were maintained in FluoroBrite DMEM (Gibco) supplemented with 10% FBS, and 2 mM L-Glutamine. Adhesion dynamics were analysed using the Focal Adhesion Analysis Server (FAAS)^93^ with the following settings: a minimum adhesion size of 20 pixels, a minimum adhesion phase length of 3 images, and a detection threshold for segmentation of 1.5 for GFP-TNS1 SG, 2.0 for WT and ΔIDR and 4.0 for SD. All the remaining settings were kept at their default values.

#### Spheroid invasion

MicroTissue 3D Petri Dishes (Sigma) were used to prepare 2% agarose moulds for spheroid formation. In each mould, 100,000 cells were seeded and incubated for 48 h. The spheroids were then embedded in a mixture of in-house rat-tail collagen (final concentration ca. 1.5 mg/ml) in 1× DMEM (Thermo Fisher Scientific) supplemented with 1% FBS, 2.7 mg/ml sodium bicarbonate (Sigma), and 20 mM HEPES (Sigma). After polymerization, the collagen gels were overlaid with cell culture medium, and the spheroids were imaged at the indicated time points using a Leica Thunder microscope equipped with a 5× objective (as described above).

#### Cell migration

Cells were seeded on fibronectin-coated µ-Slide 8 Well (Ibidi) and allowed to adhere for 3 h. Time-lapse imaging was performed over 10 h with 10 min intervals in three independent replicates using Nikon Eclipse Ti2-E equipped with 10× objective as described above. Individual cells were tracked using either TrackMate^94^ with pretrained Cellpose 2.0 segmentation^95^ or manual tracking, and the resulting tracks were visualised using a custom MATLAB script.

#### Traction force microscopy (TFM)

Cells were seeded on 10 kPa acrylamide gels with 0.2 μm fluorescent beads (FluoSpheres, Invitrogen) and left to adhere for 3 h. 1h before the beginning of the experiment, the cells were incubated in cell culture medium supplemented with 20 mM HEPES, 1 μg/ml Hoechst 33342, and 1 μg/ml WGA. Images were acquired using a spinning disk confocal microscope equipped with 40× water immersion objective as described above. Prior to imaging the relaxed gels, the cells were removed by adding 20% SDS into the media to the final concentration of 0.1%. Alignment of “before” and “after” images across all channels was performed using Fast4DReg^96^ plugin in Fiji (ImageJ). The displacement field and traction forces were calculated in MATLAB (v. R2023a) using the TFM package^97^ with subpixel correlation via image interpolation and the FTTC (Fourier Transform Traction Cytometry) method with regularisation parameter set to 0.0001, respectively. Traction forces underneath the individual cell masks were extracted using a custom R script and used for quantification.

### Statistical analysis

All experiments were performed in at least three independent biological replicates. Statistical analyses were performed using GraphPad Prism 8. Outliers were identified by the ROUT method (Q = 0.2%) and excluded from subsequent analyses. Datasets were assessed for normal distribution to determine the appropriate statistical test. Statistical tests used are specified in the corresponding figure legends.

## Acknowledgements

We thank Petra Laasola and Jenni Siivonen for outstanding technical assistance, Hellyeh Hamidi for critical reading and editing of the manuscript and all the members of the Ivaska Lab, especially Gautier Follain, for valuable comments and discussion, and Veli-Matti Leppänen for help with TNS1 purification. We further thank Peter Walsh, Chiara Giacomelli and Martin Bushell for valuable discussion. Imaging was performed at the Cell Imaging and Cytometry Core, Turku Bioscience Centre (Turku, Finland) with the support of the Finnish Advanced Microscopy Node of Euro-BioImaging Finland funded by the Research Council of Finland, FIRI grant numbers 359073, 358879. Mass spectrometry analysis was performed at the Turku Proteomics Facility, University of Turku and Åbo Akademi University. The facility is supported by Biocenter Finland. We acknowledge the computing resources provided by the CSC – IT Center for Science Ltd. (Espoo, Finland) and LUMI supercomputer, owned by the EuroHPC Joint Undertaking, hosted by CSC and the LUMI Consortium. This work was supported by the European Union’s Horizon Europe research and innovation programme under Marie Sklodowska-Curie grant agreement number 101108089 (M.D.), by the Finnish Cancer Institute (K. Albin Johansson Professorship, J.I.); a Research Council of Finland Centre of Excellence program (# 346131, J.I.); the Cancer Foundation Finland (J.I.); the Sigrid Juselius Foundation (J.I.); the Research Council of Finland’s Flagship InFLAMES (# 337530 & 357910), the Jane and Aatos Erkko Foundation (J.I.), European Research Council Advanced Grant (# 101142305; Border Control) and Research Council of Finland postdoctoral research grant (# 343239, M.R.C.). We also thank the Research Council of Finland (projects # 331349, # 336234, # 346135), the Sigrid Juselius Foundation, Helsinki Institute of Life Science (HiLIFE) Fellow Program, the Jane and Aatos Erkko Foundation, and the Lundbeck Foundation for support (I.V.).

## Author contributions

Conceptualization, M.D. and J.I.; methodology, M.D., G.E., M.R.C., I.V. and J.I.; investigation, M.D., G.E., M.R.C.; resources, M.D., G.E., M.R.C., I.V. and J.I.; writing – original draft, M.D. and J.I.; writing – review and editing, M.D., G.E., M.R.C., I.V. and J.I.; supervision, I.V. and J.I.; funding acquisition, M.D., I.V. and J.I.

## Declaration of interests

The authors declare no competing interests.

## Supplemental information

Figures S1–S7

Table S1. Document file containing the list of synthesized DNA sequences and primers used in this study.

Table S2. Excel file containing processed MS data from BioID experiment. Related to Figure 3.

Table S3. Excel file containing processed TNS1 phosphoproteomic data. Related to Figure 6.

Movie S1. Fusion of GFP-TNS1 condensates. Related to Figure 1C.

Movie S2. Fission of GFP-TNS1 condensates. Related to Figure 1D.

Movie S3. TNS1 condensate formation upon focal adhesion (FA) disassembly, related to Figure S1A.

Movie S4. GFP-TNS1 dynamics in living cells. Related to Figure S1B.

Movie S5. Representative FRAP time-lapse. Related to Figure 1E.

Movie S6. Time-lapse of control U2OS GFP-TNS1 cells. Related to Figure 6A.

Movie S7. Time-lapse of arsenite-treated U2OS GFP-TNS1 cells. Related to Figure 6A.

Movie S8. Time-lapse of U2OS GFP-TNS1 cells pre-treated with inhibitors before the arsenite treatment. Related to Figure 6E.

---

**Figure S1.**
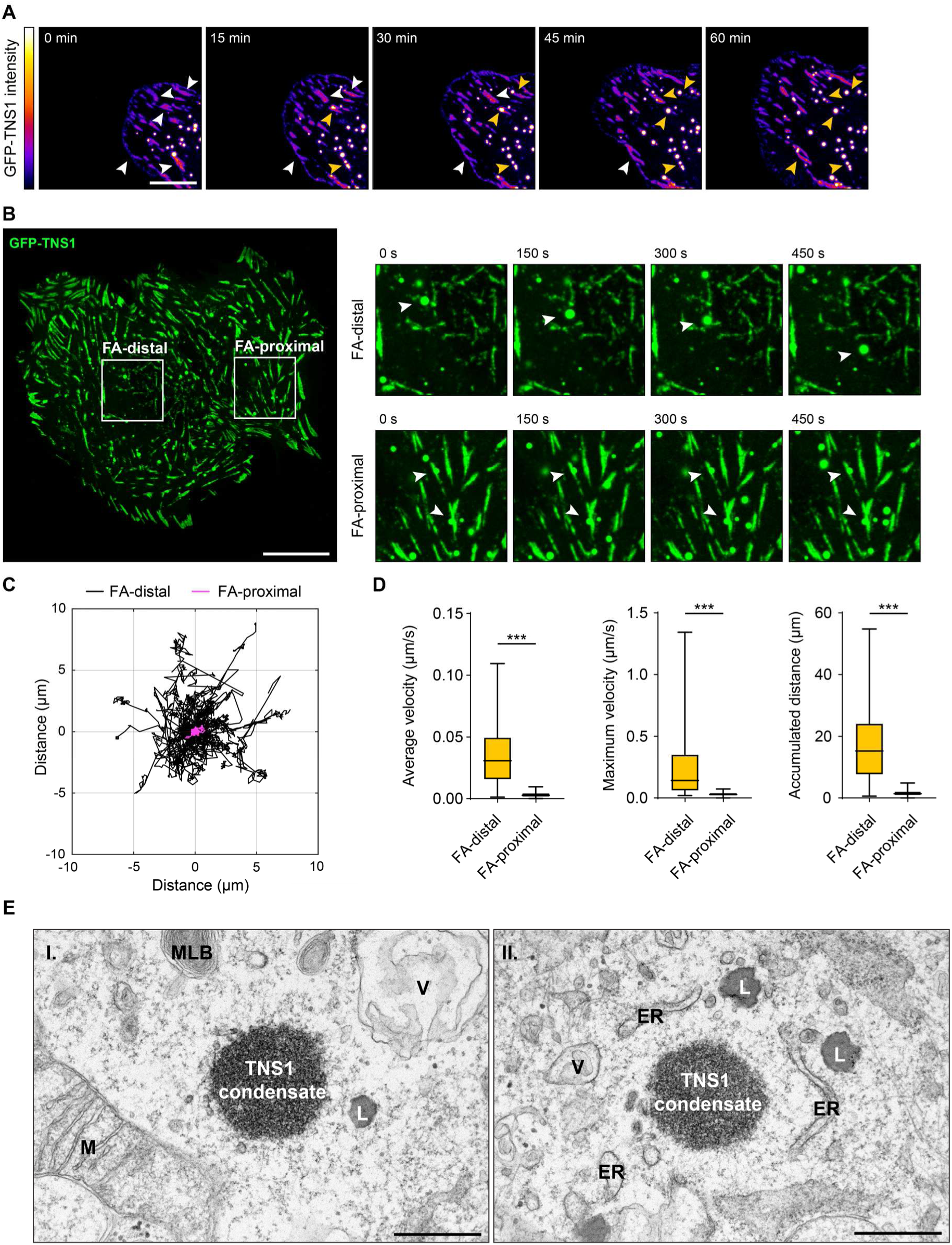
A) Representative confocal time-lapse images of TNS1 condensate formation upon focal adhesion (FA) disassembly (Movie S3). White arrows indicate FAs, yellow arrows indicate the appearance of TNS1 condensates from the respective FAs. Images acquired for 60 min at 3 min intervals. Scale bar 10 μm. B) Time-lapse images from Movie S4. Closeups illustrate behaviour of GFP-TNS1 condensates either proximal or distal to focal adhesions as indicated by arrowheads. Images acquired for 495 s with 5 s intervals. Scale bar 20 μm. C) Plot of trajectories tracking individual FA-distal (n = 74) or FA-proximal (n = 64) GFP-TNS1 condensates in U2OS cells. D) Quantification of average velocity, maximum velocity and accumulated distance of FA-distal and FA-proximal GFP-TNS1 condensates tracked in Figure S1C. Statistical analysis was performed with two-tailed Mann-Whitney U test. p < 0.001 (***). E) Annotation of subcellular structures in electron micrographs shown in Figure 1F. MLB – multilamellar body; V – vesicle; M – mitochondria; ER – endoplasmic reticulum; L – lipid droplet. Scale bar 500 nm.

**Figure S2.**
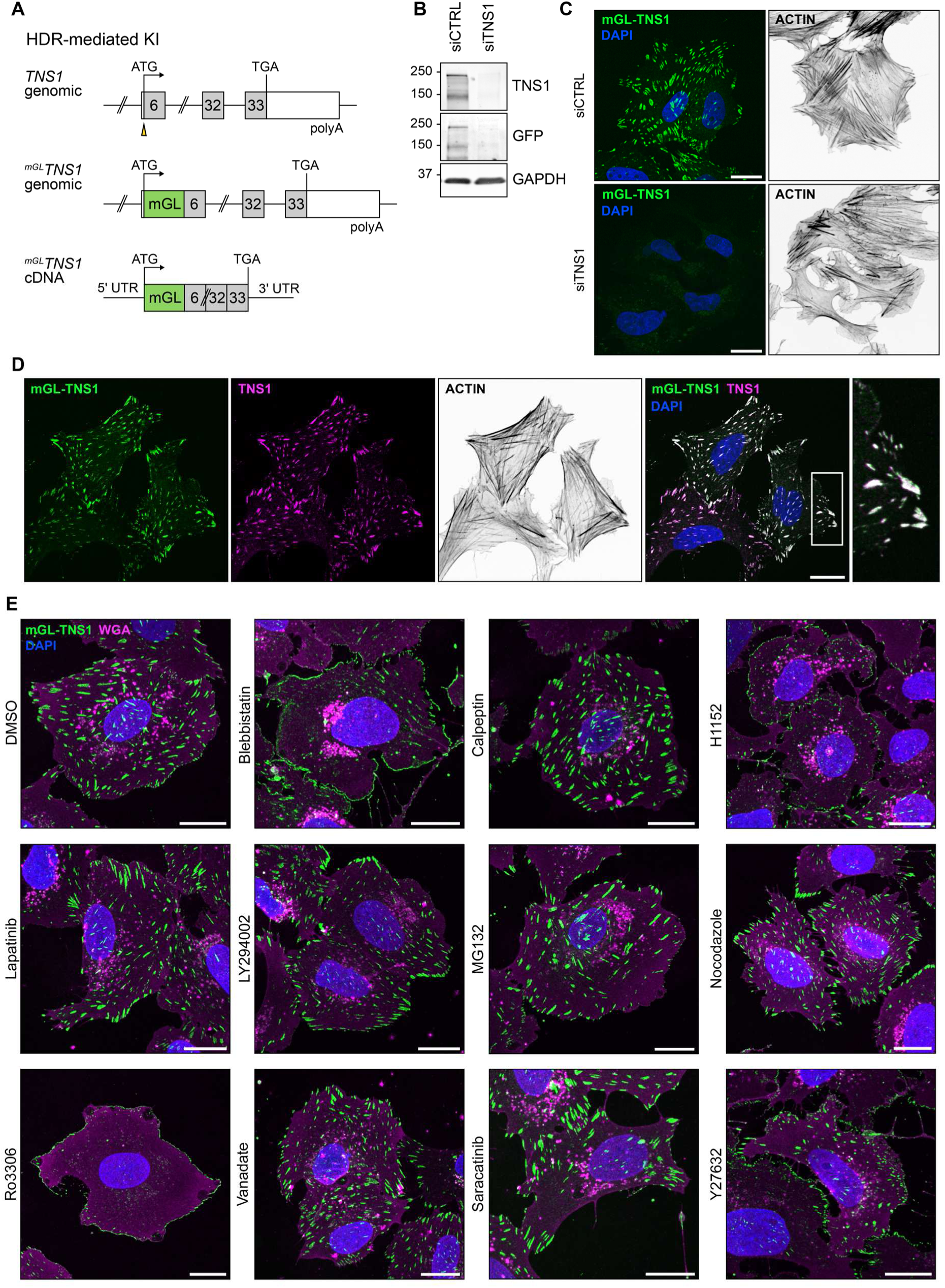
A) Schematic representation of CRISPR/Cas9-mediated knock-in of mGreenLantern (mGL) cDNA into Exon6 of the TNS1 gene. The knock-in results in production of N-terminally tagged TNS1. B) Expression of endogenous TNS1 in U2OS mGL-TNS1 cells was silenced using siRNA to confirm specificity of the knock-in. Immunoblots with anti-TNS1, anti-GFP (recognizes mGL) and anti-GAPDH antibodies are shown. C) Representative confocal images of U2OS mGL-TNS1 cells transfected with control or TNS1-specific siRNAs. Scale bars 20 μm. D) Representative confocal images of U2OS mGL-TNS1 cells stained with anti-TNS1 antibody. Actin was stained using Phalloidin. Scale bar 20 μm. E) Representative confocal images of U2OS mGL-TNS1 cells treated with the indicated compounds. Cells were stained with WGA and DAPI. Scale bars 20 μm.

**Figure S3.**
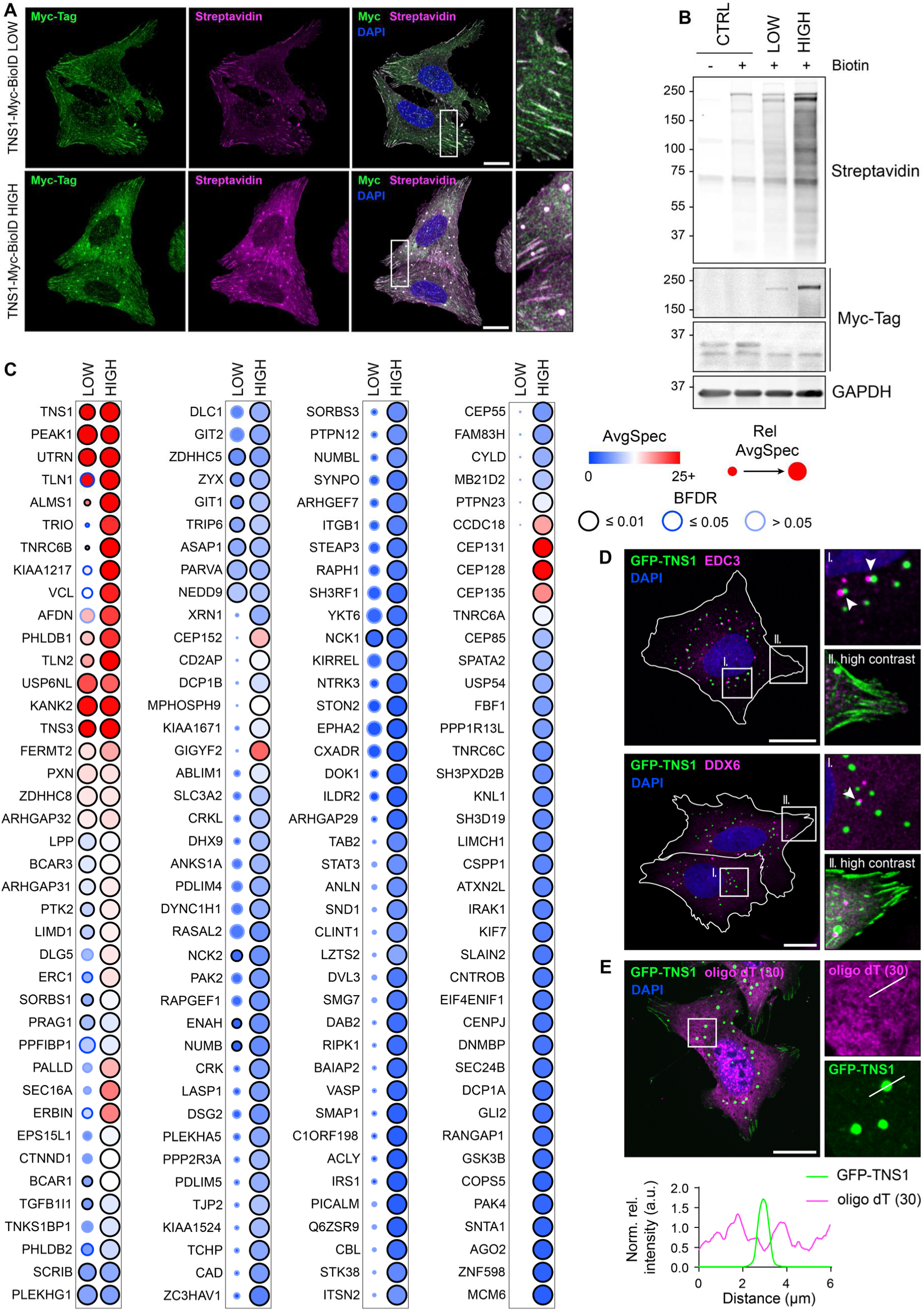
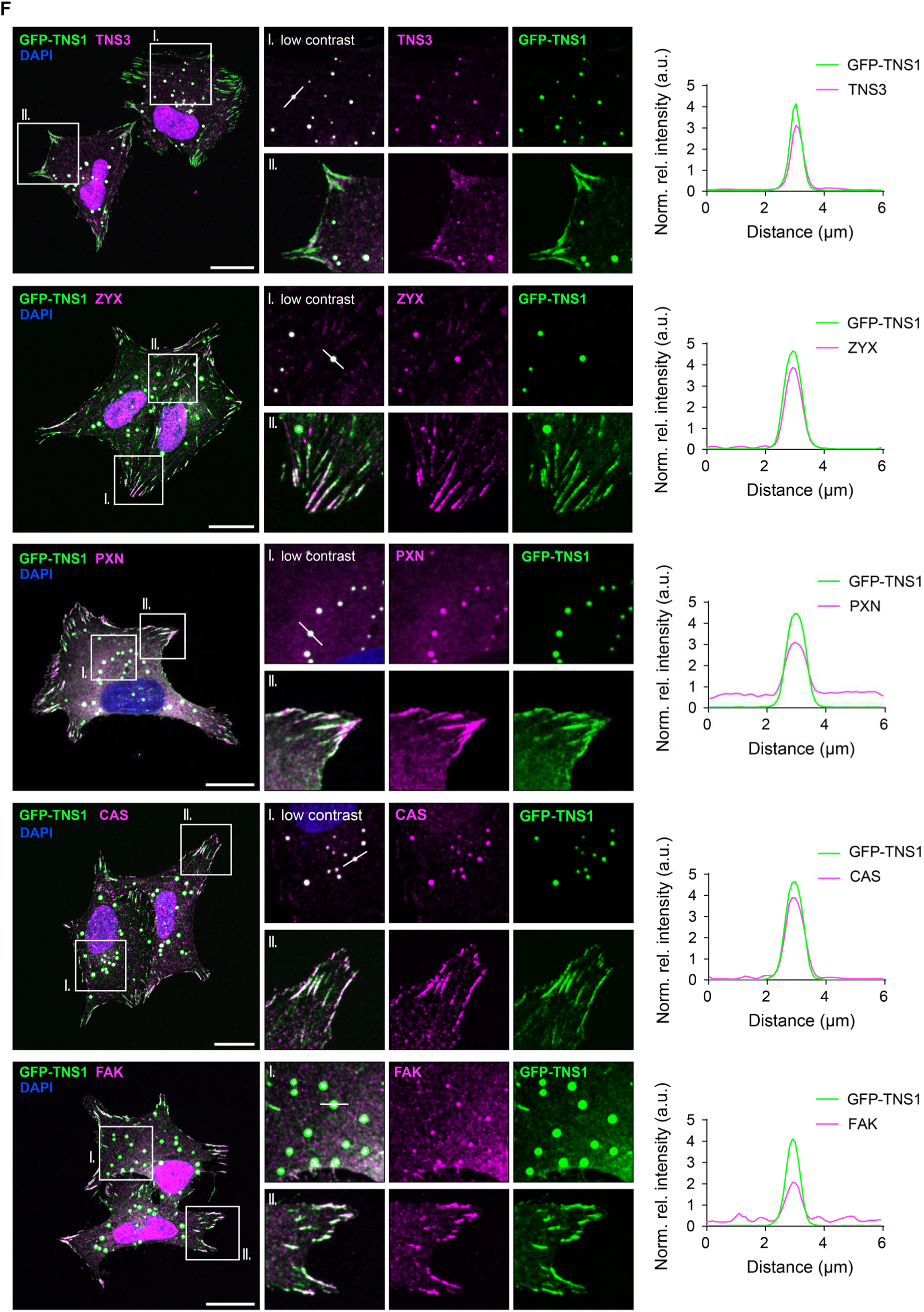
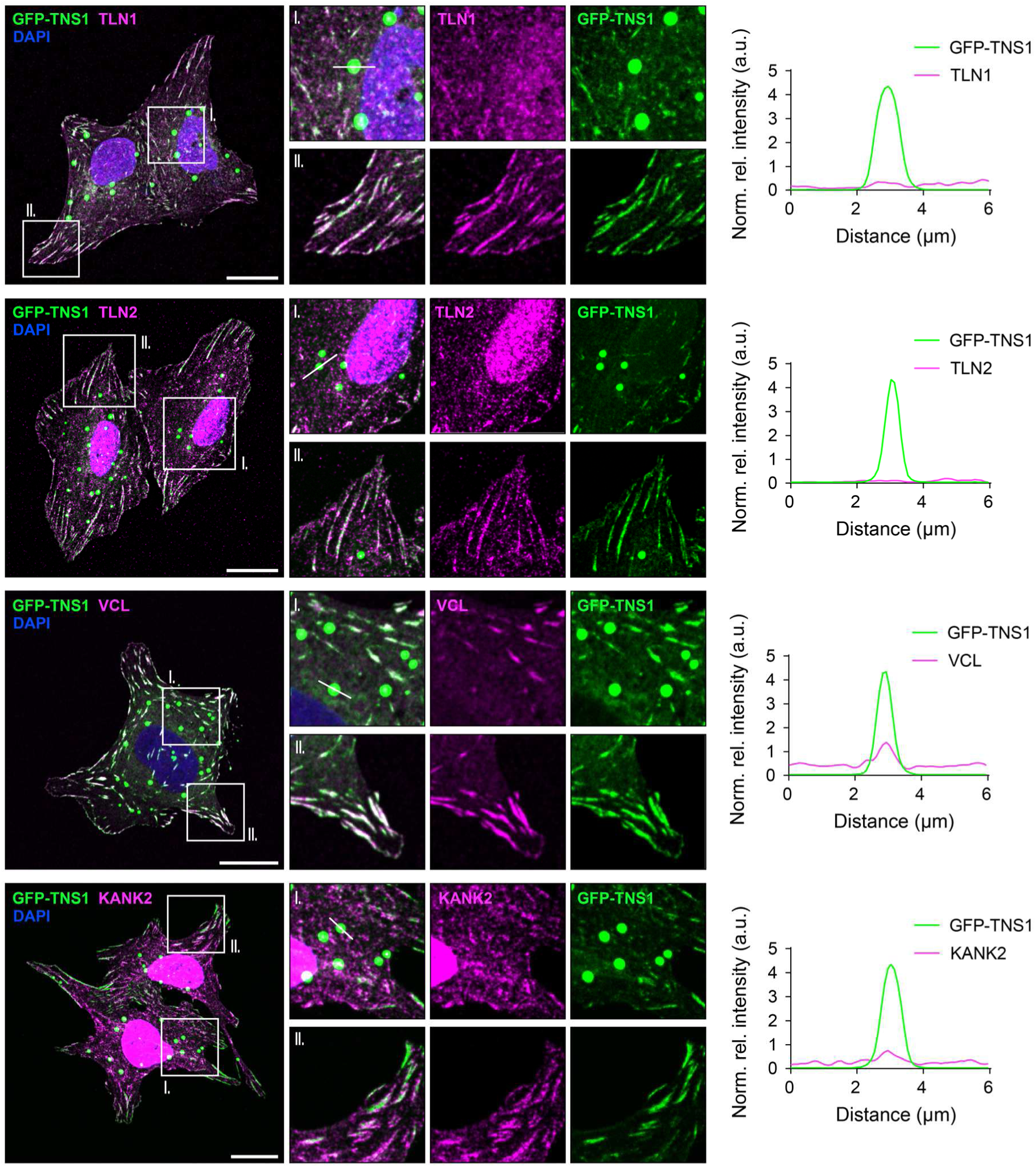
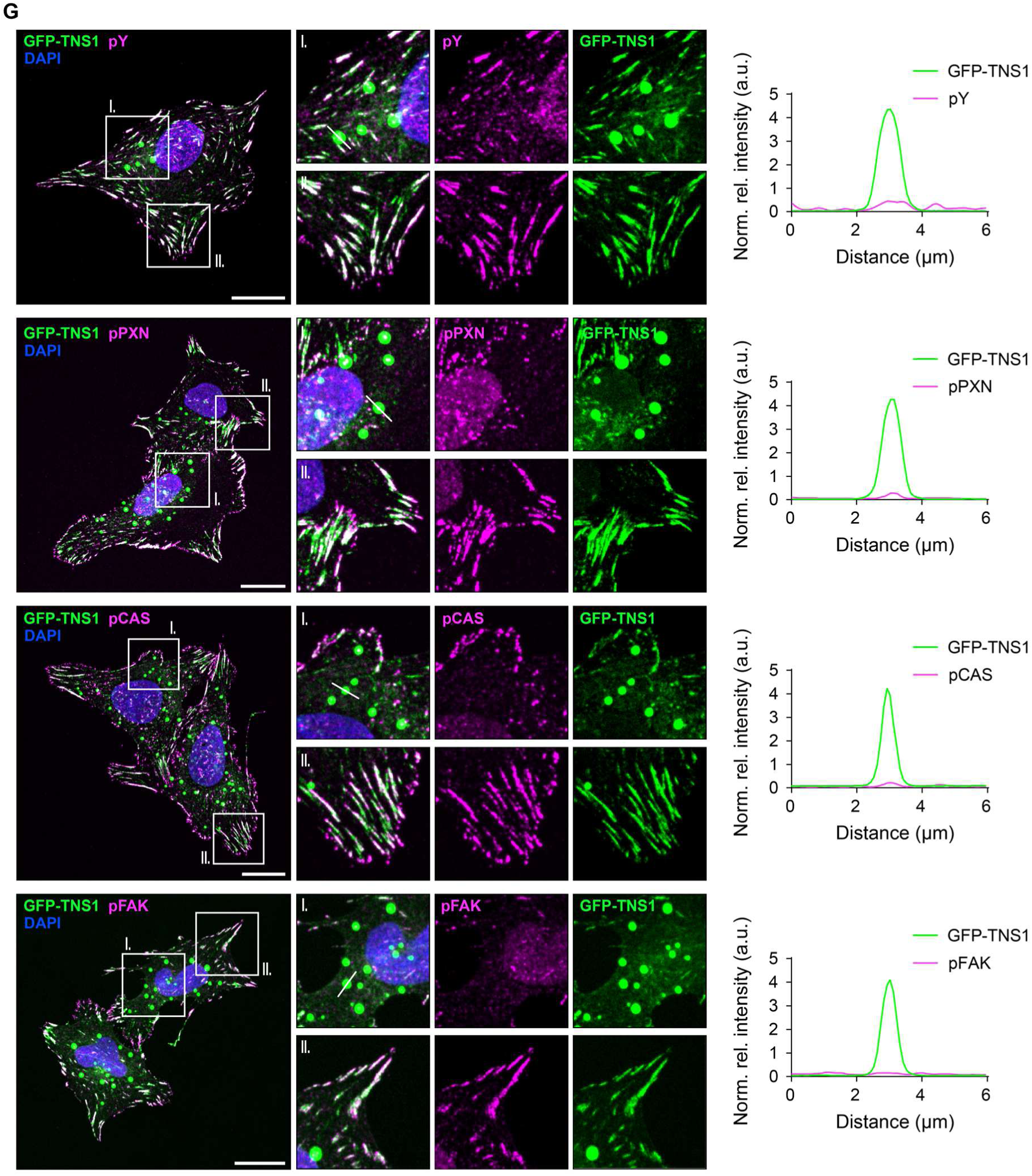
A) Representative confocal images validating correct localisation of TNS1-Myc-BioID in U2OS cells with either low or high expression of TNS1-Myc-BioID. Cells were treated with biotin for 18 h, fixed and stained with fluorescently-conjugated streptavidin, anti-Myc-Tag antibody and DAPI. Scale bars 20 μm. B) Representative immunoblots from lysates of U2OS cells expressing BioID only, and either low or high amounts of TNS1-Myc-BioID. Cells were treated with biotin for 18 h before lysis. C) High-confidence hits identified in TNS1 BioID screen. Data was visualised using ProHits Viz. Color-coding of individual spots represents the average number of unique peptides identified (AvgSpec), size of the spots reflects the relative abundance between the two conditions (Rel AvgSpec), and the color-coding of the spot outlines represents Bayesian false discovery rate (BFDR) at statistical levels as indicated. D) Representative confocal images of U2OS cells expressing GFP-TNS1. Fixed cells were stained with DAPI and either anti-EDC3 (top) or anti-DDX6 (bottom) antibodies. Closeups I. focus on association of TNS1 condensates with P-bodies (white arrowheads), closeups II. highlight the canonical TNS1 localisation in FAs. Scale bars 20 μm. E) Representative confocal image of mRNA-FISH (fluorescence *in situ* hybridisation) with oligo-dT(30) probes in U2OS cells expressing GFP-TNS1. Nuclei were visualised using DAPI staining. Profile plot (bottom) illustrates the relative intensity distribution along the white line indicated in closeups. Scale bar 20 μm. F) Representative confocal images of U2OS cells expressing GFP-TNS1 stained with either anti-CAS, anti-PXN (paxillin), anti-ZYX (zyxin), anti-TNS3 (tensin 3), anti-FAK (focal adhesion kinase), anti-TLN1 (talin 1), anti-TLN2 (talin 2), anti-KANK2 (KN motif and ankyrin repeat domains 2) or anti-VCL (vinculin) antibodies. Closeups I. focus on colocalization of GFP-TNS1 with the stained proteins in the condensates, closeups II. highlight the canonical localisation of the individual FA components in FAs. Profile plots illustrate the relative intensity distribution of individual proteins in the TNS1 condensates along the white lines indicated in closeups I. Nuclei were visualised using DAPI staining. Scale bars 20 μm. G) Representative confocal images of U2OS cells expressing GFP-TNS1 stained with either anti-pY (general phosphoTyrosine), anti-pCAS, anti-pPXN or anti-pFAK antibodies. Closeups I. focus on colocalization of GFP-TNS1 with the stained proteins in the condensates, closeups II. highlight the canonical localisation of the individual FA components in FAs. Profile plots illustrate the relative intensity distribution of individual proteins in the TNS1 condensates along the white lines indicated in closeups I. Nuclei were visualised using DAPI staining. Scale bars 20 μm.

**Figure S4.**
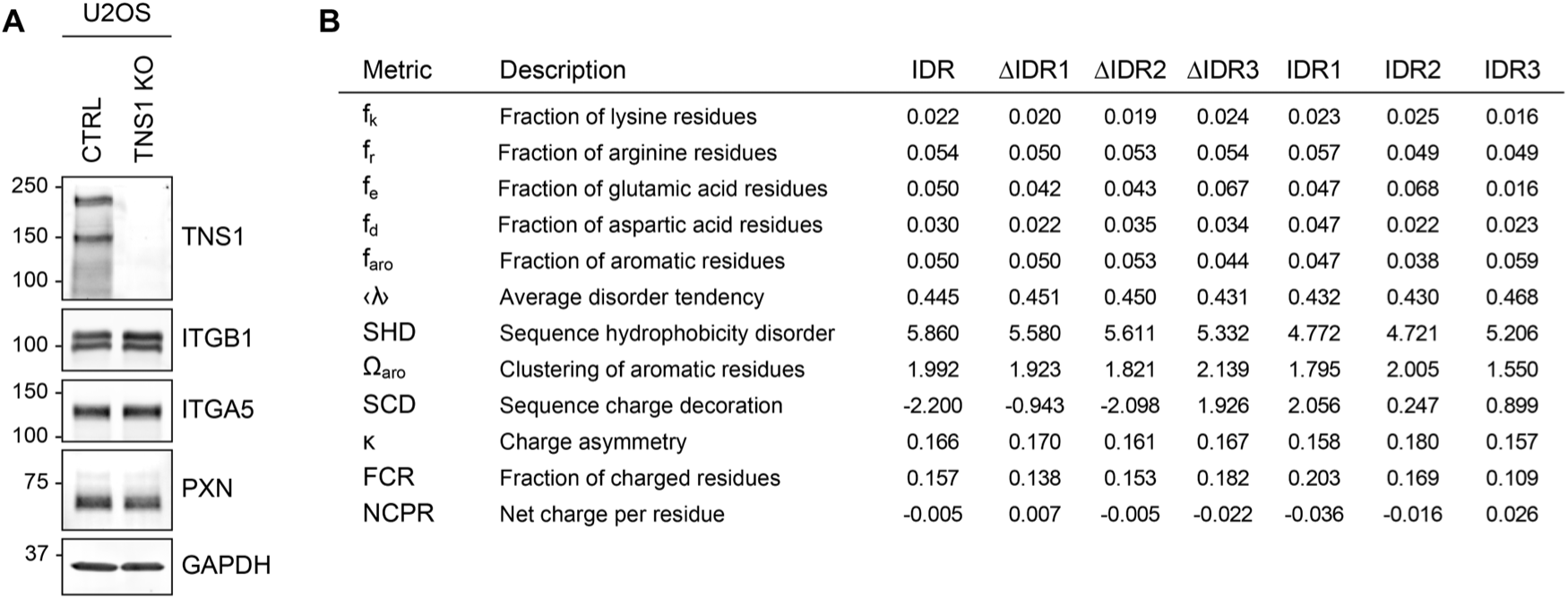
A) Representative immunoblots from lysates of control (CTRL) and TNS1 KO U2OS cells with indicated antibodies. B) Compositional analysis of the IDR sequence, its individual deletion mutants and subregions.

**Figure S5.**
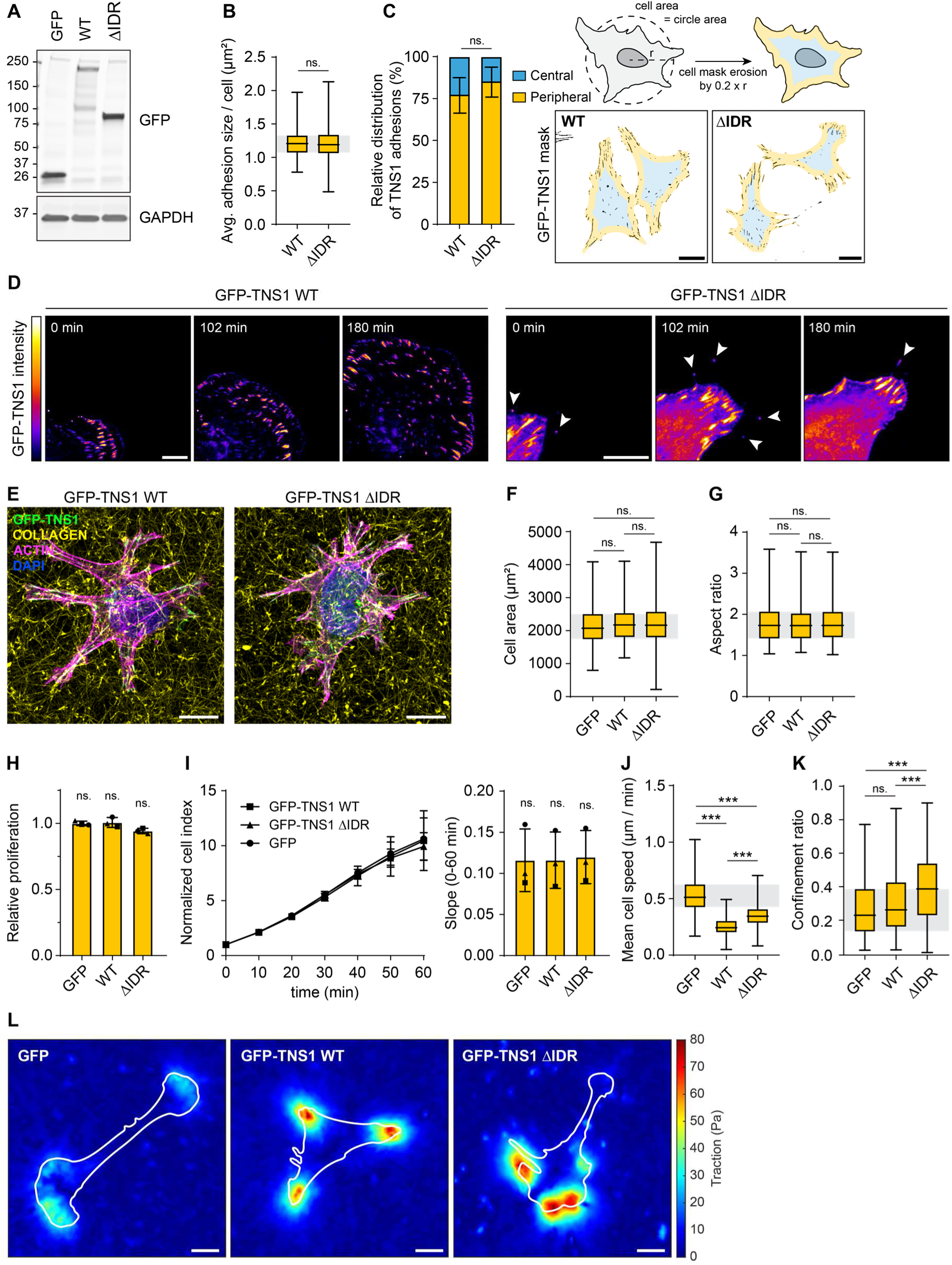
A) Representative immunoblots of U2OS TNS1 KO cells with inducible expression of either GFP or the indicated GFP-TNS1 variants. B) Quantification of the average FA size per cell from n = 155-165 cells per condition from three independent experiments. Statistical analysis was performed using two-tailed Mann-Whitney U test. ns. – not significant. C) Quantification of relative distribution of peripheral and central FAs from the indicated cell lines. Analysis was performed from n = 117-137 cells per condition by eroding the cell mask by the factor of 0.2 × radius (r) of a circle with area corresponding to the area of the cell. Representative images corresponding to central and peripheral areas with GFP-TNS1 adhesion masks are shown. Scale bars 20 μm. Statistical analysis was performed using 2-way ANOVA test. ns. – not significant. D) Representative time-lapse images of cell protrusions of indicated cell lines migrating on FN-coated coverslips. Images were acquired for 3 h at 3 min intervals. White arrowheads are indicating TNS1-containing filopodia tips. Scale bars 10 μm. E) Representative 3D projection of cells expressing indicated GFP-TNS1 variants embedded in 3D fibrillar rat tail collagen conjugated with Atto647N. The cells were stained for F-actin and DAPI was used to stain cell nuclei. Scale bars 10 μm. F) Quantification of cell area of U2OS cells expressing the indicated proteins. Quantification was performed from n = 266-294 cells per condition. Statistical analysis was performed using Kruskal-Wallis test with Dunn’s multiple comparisons test. ns. – not significant. G) Quantification of aspect ratio of U2OS cells expressing the indicated proteins. Quantification was performed from n = 262-290 cells per condition. Statistical analysis was performed using Kruskal-Wallis test with Dunn’s multiple comparisons test. ns. – not significant. H) Quantification of relative proliferation of U2OS cells expressing indicated proteins from three independent experiments. Statistical analysis was performed using Kruskal-Wallis test with Dunn’s multiple comparisons test. ns. – not significant. I) Representative plot of normalized cell index as measured by xCELLigence (left). Mean values of 3 technical replicates with the corresponding standard deviations are shown. Slopes of each condition from three independent biological replicates are shown (right). Statistical analysis was performed using Kruskal-Wallis test with Dunn’s multiple comparisons test. ns. – not significant. J) Quantification of mean cell speed of cells analysed in Figure 5F. Statistical analysis was performed using Kruskal-Wallis test with Dunn’s multiple comparisons test. p < 0.001 (***). K) Quantification of confinement ratio of cells analysed in Figure 5F. Statistical analysis was performed using Kruskal-Wallis test with Dunn’s multiple comparisons test. p < 0.001 (***); ns. – not significant. L) Representative traction map images of indicated cells seeded on 10 kPa gels. Cell outlines are shown with white line. Scale bars 20 μm.

**Figure S6.**
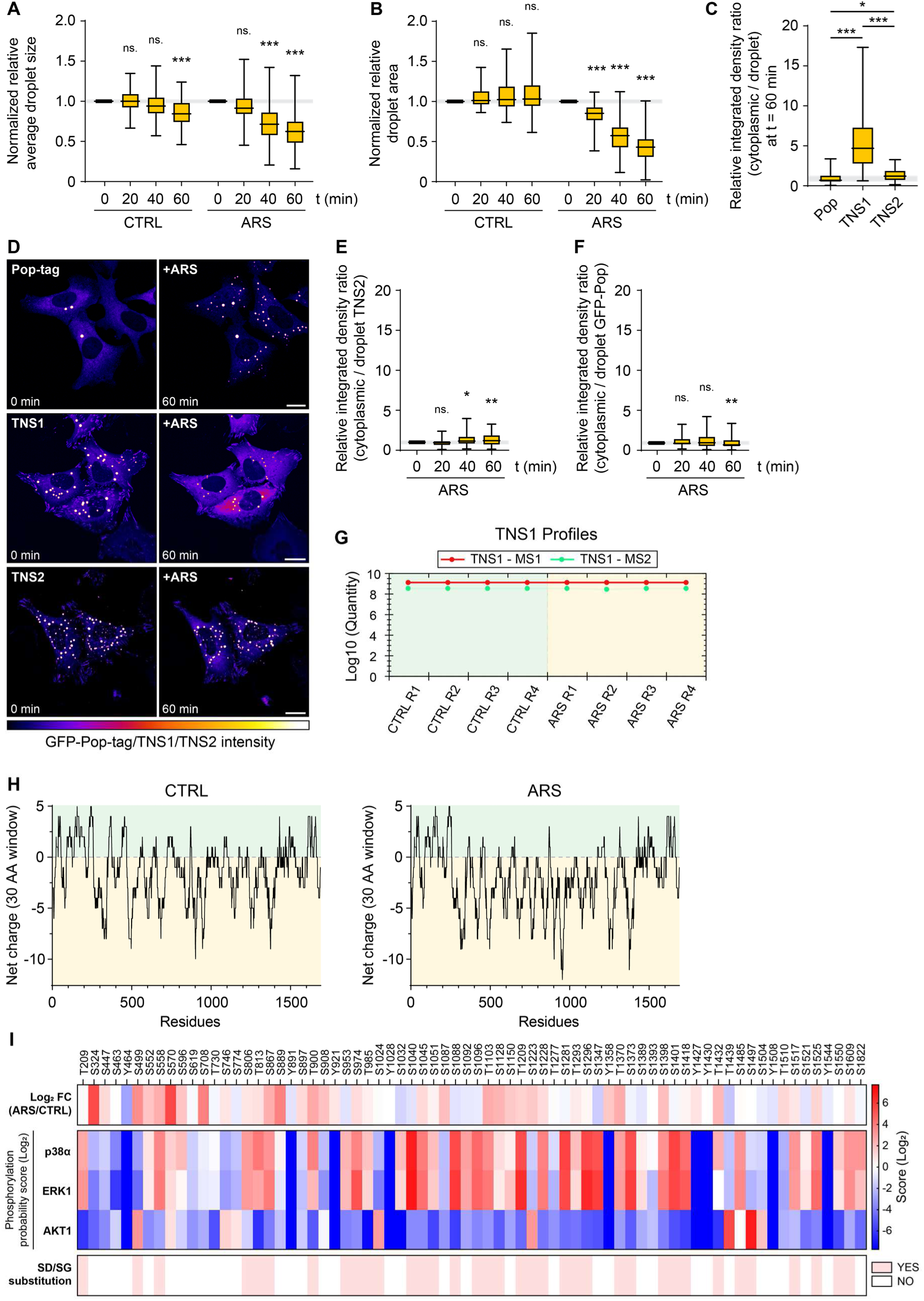
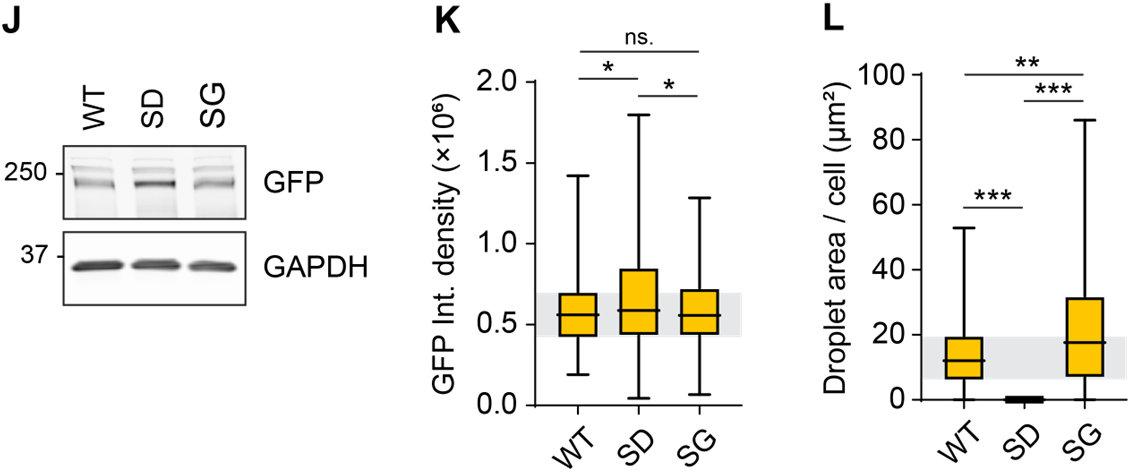
A) Quantification of normalized relative average TNS1 droplet size from n = 81-119 cells analysed in Figures 6A-B. Statistical analysis was performed using Friedman test with Dunn’s multiple comparisons test comparing the individual timepoints to t = 0 min. p < 0.001 (***); ns. – not significant. B) Quantification of normalized relative TNS1 droplet area from n = 80-117 cells analysed in Figures 6A-B. Statistical analysis was performed using Friedman test with Dunn’s multiple comparisons test comparing the individual timepoints to t = 0 min. p < 0.001 (***); ns. – not significant. C) Quantification of the ratio between the relative cytoplasmic and droplet integrated densities of GFP-Pop, GFP-TNS1 and GFP-TNS2 at t = 60 min from n = 104-115 cells per condition. Statistical analysis was performed using Kruskal-Wallis test with Dunn’s multiple comparisons test. p < 0.05 (*); p < 0.001 (***). D) Representative timelapse images of ARS-treated U2OS cells expressing either GFP-Pop, GFP-TNS1 or GFP-TNS2. Images were acquired for 60 min at 1 min intervals. Images at t = 0 min and t = 60 min are shown. Scale bars 20 μm. E) Quantification of the ratio between the relative cytoplasmic and droplet GFP-TNS2 integrated densities at the indicated timepoints from n = 112 cells. Statistical analysis was performed using Friedman test with Dunn’s multiple comparisons test comparing the individual timepoints to t = 0 min. p < 0.05 (*); p < 0.01 (**); ns. – not significant. F) Quantification of the ratio between the relative cytoplasmic and droplet GFP-Pop integrated densities at the indicated timepoints from n = 104 cells. Statistical analysis was performed using Friedman test with Dunn’s multiple comparisons test comparing the individual timepoints to t = 0 min. p < 0.01 (**); ns. – not significant. G) Plot of normalized TNS1 quantities identified by mass spectrometry. H) Cumulative net charge distribution over 30 AA window in TNS1 from CTRL (left) or ARS-treated (right) cells. I) Heatmap of phosphorylation probability score for individual phosphorylation sites by p38α, ERK1 and AKT1 kinases as predicted by Kinase Library tool^57^. Top panel represents Log_2_ fold-change of individual TNS1 phosphosites in CTRL and ARS-treated conditions. Bottom panel (SD/SG substitution) indicates phosphorylation sites selected for molecular dynamics simulations and experimental validation. J) Representative immunoblots from lysates of U2OS cells with inducible expression of the indicated GFP-TNS1 variants. K) Quantification of integrated density of cells analysed in Figures 6J-L and S6N. Statistical analysis was performed using Kruskal-Wallis test with Dunn’s multiple comparisons test. p < 0.05 (*); ns. – not significant. L) Quantification of TNS1 droplet area per cell as quantified from n = 343-358 cells per condition. Statistical analysis was performed using Kruskal-Wallis test with Dunn’s multiple comparisons test. p < 0.01 (**); p < 0.001 (***).

**Figure S7.**
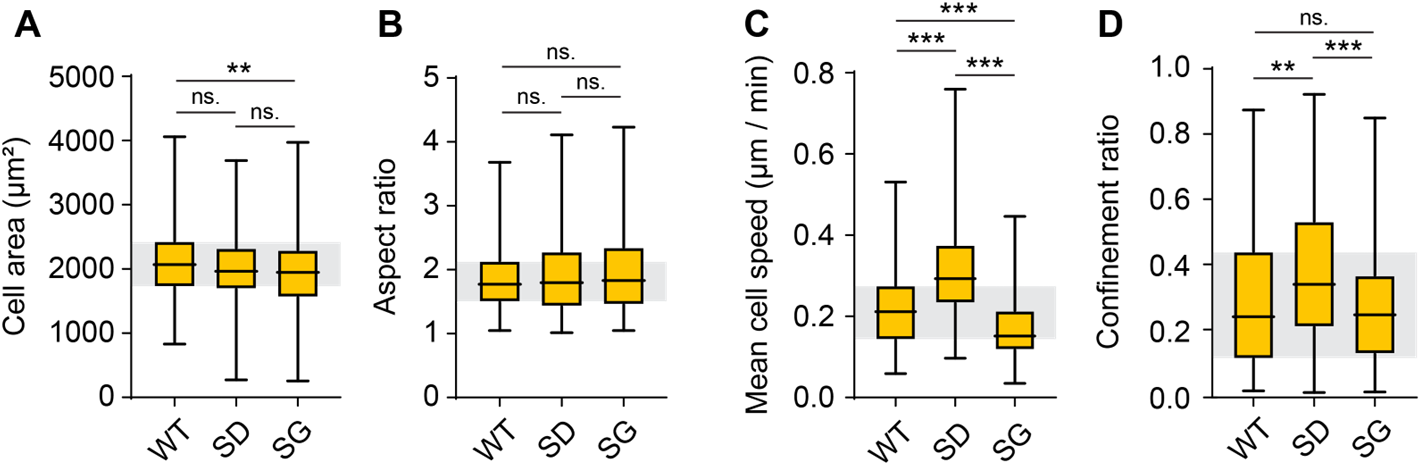
A) Quantification of cell are from n = 339-374 cells. Statistical analysis was performed using Kruskal-Wallis test with Dunn’s multiple comparisons test. p < 0.01 (**); ns. – not significant. B) Quantification of cell are from n = 331-367 cells. Statistical analysis was performed using Kruskal-Wallis test with Dunn’s multiple comparisons test. ns. – not significant. C) Quantification of mean cell speed of cells analysed in Figure 7F. Statistical analysis was performed using Kruskal-Wallis test with Dunn’s multiple comparisons test. p < 0.001 (***). D) Quantification of confinement ratio of cells analysed in Figure 7F. Statistical analysis was performed using Kruskal-Wallis test with Dunn’s multiple comparisons test. p < 0.001 (***); ns. – not significant.

## Notes

### Competing Interest Statement

The authors have declared no competing interest.

