## Supplementary material for "Adhesion-derived condensates control component availability to regulate adhesion dynamics": Table S1

**Supplementary information**

**Table S1**

**List of synthetic DNA sequences and primers used in the study**

TNS1 SD_1

GTACAAGTCCGGACTCAGATCTCGAGCTCAAGCTTACATGAGTGTGAGCCGGACCATGGACAGCTGTGAGCTGGACCTGGTGTACGTCACAGAGAGGATCATCGCTGTCTCCTTCCCCAGCACAGCCAATGAGGAGAACTTCCGGAGCAACCTCCGTGAGGTGGCGCAGATGCTCAAGTCCAAACATGGAGGCAACTACCTGCTGTTCAACCTCTCTGAGCGGAGACCTGACATCACGAAGCTCCATGCCAAGGTACTGGAATTTGGCTGGCCCGACCTCCACGATCCAGCCCTGGAGAAGATCTGCAGCATCTGTAAGGCCATGGACACATGGCTCAATGCAGACCCTCACAATGTCGTTGTTCTACACAACAAGGGAAACCGAGGCAGGATAGGAGTTGTCATCGCGGCTTACATGCACTACAGCAACATTTCTGCCAGTGCGGACCAGGCTCTGGACCAGTTTGCAATGAAGCGGTTCTATGAGGATAAGATTGTGCCCATTGGCCAGCCATCCCAAAGAAGGTACGTGCATTACTTCAGTGGCCTGCTCTCCGGCTCCATCAAAATGAACAACAAGCCCTTGTTTCTGCACCACGTGATCATGCACGGCATCCCCAACTTTGAGTCTAAAGGAGGATGTCGGCCATTTCTCCGCATCTACCAGGCCATGCAACCTGTGTACACATCTGGCATCTACAACATCCCAGGAGACAGCCAGACTAGCGTCTGCATCACCATCGAGCCAGGACTGCTCTTGAAGGGAGACATCTTGCTGAAGTGCTACCACAAGAAGTTCCGAAGCCCAGCCCGAGACGTCATCTTCCGTGTGCAGTTCCACACCTGTGCCATCCATGACCTGGGGGTTGTCTTTGGGAAGGAGGACCTTGATGATGCTTTCAAAGATGATCGATTTCCAGAGTATGGCAAAGTGGAGTTTGTATTTTCTTATGGGCCAGAGAAAATTCAAGGCATGGAGCACCTGGAGAACGGGCCGAGCGTGTCTGTGGACTATAACACCTCTGACCCCCTCATCCGCTGGGACTCCTACGACAACTTCAGTGGGCATCGAGATGACGGCATGGAGGAGGTGGTGGGACACACGCAGGGGCCACTAGATGGGAGCCTGTATGCTAAGGTGAAGAAGAAAGACGATCTGCACGGCAGCACCGGGGCTGTTAATGCCACACGTCCTACACTGTCGGCCACCCCCAACCACGTGGAACACACGCTTTCTGTGAGCAGCGACTCGGGCAACTCCACAGCCTCCACCAAGACCGACAAGACCGACGAGCCTGTCCCCGGGGCCTCCAGTGCCACTGCTGCCTTGGATCCCCAGGAGAAGCGGGAGCTGGACCGCCTGCTGAGTGGCTTTGGCTTAGAGCGAGAGAAGCAAGGCGCCATGTACCACACCCAGCACCTCAGGTCCCGCCCAGCAGGGGGCTCGGCTGTGCCCTCCTCTGGACGCCACGTTGTCCCAGCCCAGGTTCATGTCAATGGTGGGGCGTTAGCATCTGAGCGGGAGACAGACATCCTGGACGATGAATTGCCAAACCAGGATGGTCACAGTGCGGGCAGCATGGGCACACTCTCTTCTCTGGACGGGGTCACCAACACCAGTGAGGGGGGCTACCCAGAGGCCCTGTCCCCACTGACCAACGGTCTGGACAAGTCCTACCCCATGGAGCCTATGGTCAATGGAGGAGGCTACCCCTACGAGTCTGCCAGCCGGGCGGGGCCTGCCCATGCTGGCCACACGGCCCCCATGCGGCCCTCCTACTCTGCACAGGAGGGTTTAGCTGGCTACCAGAGGGAGGGGCCCCACCCAGCCTGGCCACAGCCAGTGACCACCTCCCACTATGCCCATGACCCCAGCGGTATGTTCCGCTCTCAATCCTTTTCGGAAGCTGAACCCCAGCTGCCCCCAGCTCCGGTCCGAGGGGGAAGCAGCCGGGAGGCTGTGCAAAGGGGACTGAATTCATGGCAGCAGCAGCAGCAGCAGCAGCAGCAGCCTCGCCCACCTCCACGCCAGCAGGAAAGAGCCCACTTGGAGAGTCTTGTAGCCAGCAGGCCCGATCCTCAGCCATTGGCAGAGGATCCCATCCCCAGTCTCCCTGAGTTCCCGCGAGCAGCCTCCCAGCAGGAGATTGAACAGTCCATCGAAACACTCAATATGCTGATGCTGGACCTGGAGCCAGCCTCCGCTGCTGCCCCACTACACAAGTCCCAGAGTGTCCCCGGGGCCTGGCCAGGGGCTGATCCACTCTCCTCCCAGCCCCTCTCTGGATCCTCCCGTCAGTCCCATCCACTGACCCAGTCCAGATCTGGCTATATCCCCAGTGGGCATTCGTTGGGAGATCCTGAGCCAGCCCCACGGGCCTCTCTGGAGTCTGTCCCTCCTGGCAGGTCTTACTCACCTTATGACTATCAGCCATGTTTGGCTGGGCCTAACCAGGATTTCCATTCAAAGAGCCCAGCCTCTTCCTCCTTGCCTGCCTTCCTTCCGACCACCCACGATCCTCCAGGGCCTCAGCAACCCCCAGCCTCTCTCCCTGGCCTCACTGCTCAGCCTCTGCTCGATCCAAAGGAAGCGACTTCAGACCCCTCCCGGGATCCAGAGGAGGAGCCATTGAATTTAGAAGGGCTGGTGGCCCACAGGGTAGCAGGGGTACAGGCTCGGGAGAAGCAGCCTGCAGAGCCCCCAGCCCCTCTGCGGAGGCGGGCGGCCGATGATGGACAGTATGAGAACCAGTCTCCAGAAGCCACATCCCCTCGTGATCCTGGGGTTCGCGATCCTGTCCAGTGTGTCGATCCGGAGCTGGCTCTTACCATCGCTCTCAATCCTGGAGGGCGGCCCAAAGAGCCCCATTTGCACAGCTACAAGGAGGCCTTCGAGGAGATGGAGGGAACCTCCCCGAGC

TNS1 SD_2

TGGAGGGAACCTCCCCGAGCGATCCACCACCCAGTGGGGTGCGGGATCCCCCGGGTCTGGCCAAGGATCCCCTGTCTGCTCTGGGCCTGAAACCTCACAACCCAGCGGACATCCTGTTGCACCCCACAGGAGAGCCCCGGGATTATGTGGAGTCTGTGGCACGGACAGCGGTGGCTGGACCCCGAGCTCAGGACTCTGAGCCCAAGAGCTTTAGTGCTCCAGCCACCCAGGCCTATGGCCATGAGATACCCCTGAGGAACGGGACCCTGGGTGGCTCCTTTGTCTCCCCCAGCCCCCTCTCCACCAGCAGCCCCATCCTCAGTGCTGACAGCACTTCAGTGGGGAGTTTCCCGTCGGGAGAGAGCAGTGACCAGGGTCCCCGGGATCCCACCCAGCCTCTGTTGGAGTCTGGCTTCCGCTCAGGCGATCTGGGACAGCCCGATCCATCTGCCCAGAGAAACTACCAGAGCTCTTCTCCTCTCCCGACTGTGGGCAGTAGCTACAGCAGCCCCGACTACTCACTTCAGCATTTCAGCTCCTCTCCGGAAAGCCAGGCCCGAGCTCAGTTCAGTGTGGCTGGCGTCCACACGGTGCCTGGGGATCCTCAGGCGCGCCACAGAACAGTGGGCACCAACGATCCCCCTGATCCTGGCTTCGGCCGGCGGGCCATCAATCCCAGCATGGCTGCCCCCAGCAGTCCCAGTTTGAGCCATCACCAGATGATGGGTCCACCAGGCACTGGCTTCCATGGTAGCACTGTCTCCAGCCCCCAGAGCAGTGCAGCGACCACCCCGGGGAGCCCCAGCCTGTGTCGGCACCCAGCAGGGGTCTACCAGGTTTCTGGCCTCCACAACAAAGTGGCCACCGATCCGGGGGATCCCAGCCTGGGCCGGCACCCTGGGGCTCACCAAGGCAACCTGGCCTCCGGTCTTCATAGCAATGCAATAGCCGATCCTGGAGATCCCAGCCTGGGCCGTCACCTCGGAGGGTCTGGATCTGTGGTTCCCGGCGATCCCTGCTTGGACCGGCATGTGGCCTATGGCGGCTATTCTGATCCGGAGGATCGGAGACCCACACTGTCCCGGCAGAGCAGTGCCTCTGGCTACCAGGCTCCTTCCACGCCCTCCTTCCCTGTCTCCCCTGCCTACTACCCTGGCCTGAGCAGCCCTGCCACCTCCCCGTCACCAGACTCCGCAGCCTTCCGGCAAGGGGATCCAACACCAGCCTTGCCAGAGAAGCGAAGGATGGATGTGGGAGACCGGGCAGGCAGCCTCCCCAACTATGCCACCATCAATGGGAAGGTGTCTGATCCTGTCGCCAGCGGCATGTCCGATCCCAGCGGGGGCAGCACCGTCTCCTTCTCCCACACTCTGCCCGACTTCTCCAAGTACTCCATGCCAGACAACGATCCGGAGACGCGGGCTAAAGTGAAGTTTGTCCAGGACACTTCTAAGTATTGGTACAAGCCTGAGATCTCCAGGGAGCAGGCCATCGCGCTCCTCAAGGACCAGGAGCCGGGGGCCTTCATCATCCGCGACAGTCACTCCTTCCGAGGCGCGTACGGGCTGGCCATGAAGGTGTCTGATCCACCTCCAACCATCATGCAGCAGAATAAAAAAGGAGACATGACCCATGAGCTGGTCAGGCATTTTCTGATAGAGACTGGCCCCAGAGGAGTCAAGCTCAAGGGCTGCCCCAATGAGCCAAACTTCGGATCGCTGTCTGCCCTGGTCTACCAGCACTCCATCATCCCATTGGCCCTGCCTTGCAAGCTGGTCATTCCAAACCGAGACCCCACAGATGAATCGAAAGATAGCTCCGGCCCTGCCAACTCAACTGCAGACCTGCTGAAACAAGGGGCAGCCTGCAATGTGCTCTTCATCAACTCTGTGGACATGGAGTCACTCACTGGGCCACAGGCCATCTCTAAAGCCACATCTGAGACGTTGGCTGCAGACCCCACACCAGCTGCCACCATCGTTCACTTCAAAGTCTCTGCCCAGGGAATCACTCTGACTGACAACCAGAGAAAGCTCTTTTTCAGACGCCACTACCCTCTCAACACTGTCACCTTCTGTGACCTGGATCCACAGGAAAGAAAGTGGATGAAAACAGAGGGTGGTGCCCCTGCTAAGCTCTTCGGCTTCGTGGCCCGGAAGCAGGGCAGCACCACGGACAACGCCTGCCACCTCTTTGCTGAGCTTGACCCCAACCAGCCGGCCTCTGCCATCGTCAACTTCGTCTCCAAGGTCATGCTGAATGCCGGCCAAAAGAGATGAGTCGACGGTACCGCGGGCCCGGGAT

TNS1 SG_1

GTACAAGTCCGGACTCAGATCTCGAGCTCAAGCTTACATGAGTGTGAGCCGGACCATGGACAGCTGTGAGCTGGACCTGGTGTACGTCACAGAGAGGATCATCGCTGTCTCCTTCCCCAGCACAGCCAATGAGGAGAACTTCCGGAGCAACCTCCGTGAGGTGGCGCAGATGCTCAAGTCCAAACATGGAGGCAACTACCTGCTGTTCAACCTCTCTGAGCGGAGACCTGACATCACGAAGCTCCATGCCAAGGTACTGGAATTTGGCTGGCCCGACCTCCACGGACCAGCCCTGGAGAAGATCTGCAGCATCTGTAAGGCCATGGACACATGGCTCAATGCAGACCCTCACAATGTCGTTGTTCTACACAACAAGGGAAACCGAGGCAGGATAGGAGTTGTCATCGCGGCTTACATGCACTACAGCAACATTTCTGCCAGTGCGGACCAGGCTCTGGACCAGTTTGCAATGAAGCGGTTCTATGAGGATAAGATTGTGCCCATTGGCCAGCCATCCCAAAGAAGGTACGTGCATTACTTCAGTGGCCTGCTCTCCGGCTCCATCAAAATGAACAACAAGCCCTTGTTTCTGCACCACGTGATCATGCACGGCATCCCCAACTTTGAGTCTAAAGGAGGATGTCGGCCATTTCTCCGCATCTACCAGGCCATGCAACCTGTGTACACATCTGGCATCTACAACATCCCAGGAGACAGCCAGACTAGCGTCTGCATCACCATCGAGCCAGGACTGCTCTTGAAGGGAGACATCTTGCTGAAGTGCTACCACAAGAAGTTCCGAAGCCCAGCCCGAGACGTCATCTTCCGTGTGCAGTTCCACACCTGTGCCATCCATGACCTGGGGGTTGTCTTTGGGAAGGAGGACCTTGATGATGCTTTCAAAGATGATCGATTTCCAGAGTATGGCAAAGTGGAGTTTGTATTTTCTTATGGGCCAGAGAAAATTCAAGGCATGGAGCACCTGGAGAACGGGCCGAGCGTGTCTGTGGACTATAACACCTCTGACCCCCTCATCCGCTGGGACTCCTACGACAACTTCAGTGGGCATCGAGATGACGGCATGGAGGAGGTGGTGGGACACACGCAGGGGCCACTAGATGGGAGCCTGTATGCTAAGGTGAAGAAGAAAGACGGACTGCACGGCAGCACCGGGGCTGTTAATGCCACACGTCCTACACTGTCGGCCACCCCCAACCACGTGGAACACACGCTTTCTGTGAGCAGCGACTCGGGCAACTCCACAGCCTCCACCAAGACCGACAAGACCGACGAGCCTGTCCCCGGGGCCTCCAGTGCCACTGCTGCCTTGGGACCCCAGGAGAAGCGGGAGCTGGACCGCCTGCTGAGTGGCTTTGGCTTAGAGCGAGAGAAGCAAGGCGCCATGTACCACACCCAGCACCTCAGGTCCCGCCCAGCAGGGGGCTCGGCTGTGCCCTCCTCTGGACGCCACGTTGTCCCAGCCCAGGTTCATGTCAATGGTGGGGCGTTAGCATCTGAGCGGGAGACAGACATCCTGGACGATGAATTGCCAAACCAGGATGGTCACAGTGCGGGCAGCATGGGCACACTCTCTTCTCTGGACGGGGTCACCAACACCAGTGAGGGGGGCTACCCAGAGGCCCTGTCCCCACTGACCAACGGTCTGGACAAGTCCTACCCCATGGAGCCTATGGTCAATGGAGGAGGCTACCCCTACGAGTCTGCCAGCCGGGCGGGGCCTGCCCATGCTGGCCACACGGCCCCCATGCGGCCCTCCTACTCTGCACAGGAGGGTTTAGCTGGCTACCAGAGGGAGGGGCCCCACCCAGCCTGGCCACAGCCAGTGACCACCTCCCACTATGCCCATGACCCCAGCGGTATGTTCCGCTCTCAATCCTTTTCGGAAGCTGAACCCCAGCTGCCCCCAGCTCCGGTCCGAGGGGGAAGCAGCCGGGAGGCTGTGCAAAGGGGACTGAATTCATGGCAGCAGCAGCAGCAGCAGCAGCAGCAGCCTCGCCCACCTCCACGCCAGCAGGAAAGAGCCCACTTGGAGAGTCTTGTAGCCAGCAGGCCCGGACCTCAGCCATTGGCAGAGGGACCCATCCCCAGTCTCCCTGAGTTCCCGCGAGCAGCCTCCCAGCAGGAGATTGAACAGTCCATCGAAACACTCAATATGCTGATGCTGGACCTGGAGCCAGCCTCCGCTGCTGCCCCACTACACAAGTCCCAGAGTGTCCCCGGGGCCTGGCCAGGGGCTGGACCACTCTCCTCCCAGCCCCTCTCTGGATCCTCCCGTCAGTCCCATCCACTGACCCAGTCCAGATCTGGCTATATCCCCAGTGGGCATTCGTTGGGAGGACCTGAGCCAGCCCCACGGGCCTCTCTGGAGTCTGTCCCTCCTGGCAGGTCTTACTCACCTTATGACTATCAGCCATGTTTGGCTGGGCCTAACCAGGATTTCCATTCAAAGAGCCCAGCCTCTTCCTCCTTGCCTGCCTTCCTTCCGACCACCCACGGACCTCCAGGGCCTCAGCAACCCCCAGCCTCTCTCCCTGGCCTCACTGCTCAGCCTCTGCTCGGACCAAAGGAAGCGACTTCAGACCCCTCCCGGGGACCAGAGGAGGAGCCATTGAATTTAGAAGGGCTGGTGGCCCACAGGGTAGCAGGGGTACAGGCTCGGGAGAAGCAGCCTGCAGAGCCCCCAGCCCCTCTGCGGAGGCGGGCGGCCGGAGATGGACAGTATGAGAACCAGTCTCCAGAAGCCACATCCCCTCGTGGACCTGGGGTTCGCGGACCTGTCCAGTGTGTCGGACCGGAGCTGGCTCTTACCATCGCTCTCAATCCTGGAGGGCGGCCCAAAGAGCCCCATTTGCACAGCTACAAGGAGGCCTTCGAGGAGATGGAGGGAACCTCCCCGAGC

TNS1 SG_2

TGGAGGGAACCTCCCCGAGCGGACCACCACCCAGTGGGGTGCGGGGACCCCCGGGTCTGGCCAAGGGACCCCTGTCTGCTCTGGGCCTGAAACCTCACAACCCAGCGGACATCCTGTTGCACCCCACAGGAGAGCCCCGGGGATATGTGGAGTCTGTGGCACGGACAGCGGTGGCTGGACCCCGAGCTCAGGACTCTGAGCCCAAGAGCTTTAGTGCTCCAGCCACCCAGGCCTATGGCCATGAGATACCCCTGAGGAACGGGACCCTGGGTGGCTCCTTTGTCTCCCCCAGCCCCCTCTCCACCAGCAGCCCCATCCTCAGTGCTGACAGCACTTCAGTGGGGAGTTTCCCGTCGGGAGAGAGCAGTGACCAGGGTCCCCGGGGACCCACCCAGCCTCTGTTGGAGTCTGGCTTCCGCTCAGGCGGACTGGGACAGCCCGGACCATCTGCCCAGAGAAACTACCAGAGCTCTTCTCCTCTCCCGACTGTGGGCAGTAGCTACAGCAGCCCCGACTACTCACTTCAGCATTTCAGCTCCTCTCCGGAAAGCCAGGCCCGAGCTCAGTTCAGTGTGGCTGGCGTCCACACGGTGCCTGGGGGACCTCAGGCGCGCCACAGAACAGTGGGCACCAACGGACCCCCTGGACCTGGCTTCGGCCGGCGGGCCATCAATCCCAGCATGGCTGCCCCCAGCAGTCCCAGTTTGAGCCATCACCAGATGATGGGTCCACCAGGCACTGGCTTCCATGGTAGCACTGTCTCCAGCCCCCAGAGCAGTGCAGCGACCACCCCGGGGAGCCCCAGCCTGTGTCGGCACCCAGCAGGGGTCTACCAGGTTTCTGGCCTCCACAACAAAGTGGCCACCGGACCGGGGGGACCCAGCCTGGGCCGGCACCCTGGGGCTCACCAAGGCAACCTGGCCTCCGGTCTTCATAGCAATGCAATAGCCGGACCTGGAGGACCCAGCCTGGGCCGTCACCTCGGAGGGTCTGGATCTGTGGTTCCCGGCGGACCCTGCTTGGACCGGCATGTGGCCTATGGCGGCTATTCTGGACCGGAGGATCGGAGACCCACACTGTCCCGGCAGAGCAGTGCCTCTGGCTACCAGGCTCCTTCCACGCCCTCCTTCCCTGTCTCCCCTGCCTACTACCCTGGCCTGAGCAGCCCTGCCACCTCCCCGTCACCAGACTCCGCAGCCTTCCGGCAAGGGGGACCAACACCAGCCTTGCCAGAGAAGCGAAGGATGGGAGTGGGAGACCGGGCAGGCAGCCTCCCCAACTATGCCACCATCAATGGGAAGGTGTCTGGACCTGTCGCCAGCGGCATGTCCGGACCCAGCGGGGGCAGCACCGTCTCCTTCTCCCACACTCTGCCCGACTTCTCCAAGTACTCCATGCCAGACAACGGACCGGAGACGCGGGCTAAAGTGAAGTTTGTCCAGGACACTTCTAAGTATTGGTACAAGCCTGAGATCTCCAGGGAGCAGGCCATCGCGCTCCTCAAGGACCAGGAGCCGGGGGCCTTCATCATCCGCGACAGTCACTCCTTCCGAGGCGCGTACGGGCTGGCCATGAAGGTGTCTGGACCACCTCCAACCATCATGCAGCAGAATAAAAAAGGAGACATGACCCATGAGCTGGTCAGGCATTTTCTGATAGAGACTGGCCCCAGAGGAGTCAAGCTCAAGGGCTGCCCCAATGAGCCAAACTTCGGATCGCTGTCTGCCCTGGTCTACCAGCACTCCATCATCCCATTGGCCCTGCCTTGCAAGCTGGTCATTCCAAACCGAGACCCCACAGATGAATCGAAAGATAGCTCCGGCCCTGCCAACTCAACTGCAGACCTGCTGAAACAAGGGGCAGCCTGCAATGTGCTCTTCATCAACTCTGTGGACATGGAGTCACTCACTGGGCCACAGGCCATCTCTAAAGCCACATCTGAGACGTTGGCTGCAGACCCCACACCAGCTGCCACCATCGTTCACTTCAAAGTCTCTGCCCAGGGAATCACTCTGACTGACAACCAGAGAAAGCTCTTTTTCAGACGCCACTACCCTCTCAACACTGTCACCTTCTGTGACCTGGATCCACAGGAAAGAAAGTGGATGAAAACAGAGGGTGGTGCCCCTGCTAAGCTCTTCGGCTTCGTGGCCCGGAAGCAGGGCAGCACCACGGACAACGCCTGCCACCTCTTTGCTGAGCTTGACCCCAACCAGCCGGCCTCTGCCATCGTCAACTTCGTCTCCAAGGTCATGCTGAATGCCGGCCAAAAGAGATGAGTCGACGGTACCGCGGGCCCGGGAT

mGL-TNS1 HDR template sequence

TACATGAGTGTGAGCCGGACCATGGCCCCTCCCTTCCTCACCCAGTGCCCAGTGGAGAGGATGGCCAGCCCAGGGGCCTCCCCACCTCTCCGGGAAACGCATGGTGCTGGCGGCTGCCCGCCACTGCAGCAACAAGGGCTCTGTGCCCTCTTGTTTTCTGGGCCCTTAGAAATCAGGGAGGCTGCTGCCAAGAAGCAGCTCTGGCAGGCAGTGGGAGCTGCCAGAGGTGGGGAGAAGGCCAAGGAACAAAGGGCAGGAGTTGGCCTGCCTCCGGGAGGGCTGGAATGTGTGCTGTGGTCTGGCTTACCTGGAACCCAGGTGAACGTGTCACCTGAGATGCCCCCTCCCCTGAGAGGTTTCCCGCCCACTCCACAGCTCGAGAGCTAGGGGCGGGCAGTGGGAAGTCTCCGTTCCAGGGCTGGAGAGCGCCCAGCGCAGGCTCTCCTTCCTCTTCCAGAGCAGTCTGGGGCCTTTCGGGAGACTTCTGCCTGGCCAGGCAGCCAGTGTTGGGAGCCAGAGCTCCTGGGTTCCTGGGCCAGTCTGGCCCTGGCTCTGAGCTCTGGAGTCACAGGTTGAAGGCAGTGGCCAAAGGACTGATGGGAACCAAGAATTGGTGTCATCACAGTTTGGTTCTGAAATGGCCTGAGGTCCTTGCCAGCCCTACCCAGCTTGTTCTCGCCGGCACTCCCAAGCCGGCCACATTGGCCTTGCCAGAGGAAGCCTGTGGCCTTGGCCTTGGGTTGAAGAGCAGCCAAGCACCCTGGGCTGCATATGGGGGTCACACAGGGTGTGTATGGGGTGCAGCTATCTGGGGCCTTCCCTGTGGGCTGGATTCTGATATGGACCCTCCGCTGCCTGCTGTCTCCCTAGAAACATGGTGAGCAAGGGCGAGGAGCTGTTCACCGGGGTGGTGCCCATCCTGGTCGAGCTGGACGGCGACGTAAACGGCCACAAGTTCAGCGTCCGCGGCGAGGGCGAGGGCGATGCCACCAACGGCAAGCTGACCCTGAAGTTCATCTGCACCACCGGCAAGCTGCCCGTGCCCTGGCCCACCCTCGTGACCACCTTAGGCTACGGCGTGGCCTGCTTCGCCCGCTACCCCGACCACATGAAGCAGCACGACTTCTTCAAGTCCGCCATGCCCGAAGGCTACGTCCAGGAGCGCACCATCTCTTTCAAGGACGACGGTACCTACAAGACCCGCGCCGAGGTGAAGTTCGAGGGCGACACCCTGGTGAACCGCATCGTGCTGAAGGGCATCGACTTCAAGGAGGACGGCAACATCCTGGGGCACAAGCTGGAGTACAACTTCAACAGCCACAAGGTCTATATCACGGCCGACAAGCAGAAGAACGGCATCAAGGCTAACTTCAAGACCCGCCACAACGTTGAGGACGGCGGCGTGCAGCTCGCCGACCACTACCAGCAGAACACCCCCATCGGCGACGGCCCCGTGCTGCTGCCCGACAACCACTACCTGAGCCATCAGTCCAAACTGAGCAAAGACCCCAACGAGAAGCGCGATCACATGGTCCTGAAGGAGAGGGTGACCGCCGCCGGGATTACACATGACATGGACGAGCTGTACAAGGGAGGTGGTAGTGGTGGAGGAAGTGGTGGAGGTATGTCTGTATCTAGAACTATGGAGGACAGCTGTGAGCTGGACCTGGTGTACGTCACAGAGAGGATCATCGCTGTCTCCTTCCCCAGCACAGCCAATGAGGAGAACTTCCGGAGCAACCTCCGTGAGGTGGCGCAGATGCTCAAGTCCAAACATGGAGGCAACTACCTGGTGAGGATGGATCCTCGTGCCCACTGTCCTGTGCCCTCTATGCTTCCATCTGCTCACCACCTCTACATGACTGGTATCCCTGCCACCAGCCCCTGTGCCAGCATGCCCTCTTCTCTTTGTTTATCTGCCTTTCCCCTCAGGCAGCTCTCGTGCCTGAGTCTTCAATCACTCTCCAGCCCAGCTCTCCTCTAAGGAGTGTGCTGCAGCTGGAATGTAGCATCTGGAACTCTGTGTGTGCATGTGTGTGCTGGCCAGGAGAAGAGAGTACCAATAAGAGTTCAGGTTATTGAGCGCTTGCTACTAGCCAGGCCCTGTTCCAACTGCTTTCCACGCACTTAACCTTTGCAGCAGCCCTAGCAAGCAGGCACTTGCCAGAGCACAGAGAGGTGGGGTTATTTGCCTAAGGTTGCAGGGTCAGGAAGTGGCAGAGCTGGCACAGGGTCGGTGCCTCTGAGAAGTCTGAGGGCAGAGTCTTGGCAACATGTGTCTTAAGGGACCCTGGGTTGCTGGCACCTGGCTCGGCCTGGCCCAGAGGAGTCACCTTGGCACACGGGAGAGAGGTGCTGGCGGTGGTCTCCACTAACGCAGAAAGGCTGATCCGTCCTTGGCCCCAGAGGAGGGCTCAGGTCCATTTTAGAGGGAGGTGCCTGTGGACTGGACCCACAGATCTGATCTGTACAACCTCAGGTGGCTGTCCCCACATGAGTGTGAGCCGGACCATGGAT

| **Primer** | **Sequence** |
| --- | --- |
| TNS1 dPTP F | TCAGATCTCGAGCTCAAGCTTACATGAACAACAAGCCCTTGTTTCT |
| TNS1 dPTP R | AGAAACAAGGGCTTGTTGTTCATGTAAGCTTGAGCTCGAGATCTGA |
| TNS1 dC2 F | GCTCTCCGGCTCCATCAAACCAGAGAAAATTCAAGGCATGGAG |
| TNS1 dC2 R | CTCCATGCCTTGAATTTTCTCTGGTTTGATGGAGCCGGAGAGC |
| TNS1 dIDR F | GCTGGGACTCCTACGACGCTAAAGTGAAGTTTGTCCAGGAC |
| TNS1 dIDR R | GTCCTGGACAAACTTCACTTTAGCGTCGTAGGAGTCCCAGC |
| TNS1 dIDR1 F | GCTGGGACTCCTACGACACACTCAATATGCTGATGCTGG |
| TNS1 dIDR1 R | CCAGCATCAGCATATTGAGTGTGTCGTAGGAGTCCCAGC |
| TNS1 dIDR2 F | CAGGAGATTGAACAGTCCATCGAACCCATCCTCAGTGCTGAC |
| TNS1 dIDR2 R | GTCAGCACTGAGGATGGGTTCGATGGACTGTTCAATCTCCTG |
| TNS1 dIDR3 F | CCCTCTCCACCAGCAGCGCTAAAGTGAAGTTTGTCCAGGACACT |
| TNS1 dIDR3 R | AGTGTCCTGGACAAACTTCACTTTAGCGCTGCTGGTGGAGAGGG |
| TNS1 dSH2 F | GTTTGTCCAGGACACTTCTAAGTATCGAGACCCCACAGATGAATC |
| TNS1 dSH2 R | GATTCATCTGTGGGGTCTCGATACTTAGAAGTGTCCTGGACAAAC |
| TNS1 dPTB F | CAGACCTGCTGAAACAAGGGTGAGTCGACGGTACCGC |
| TNS1 dPTB R | GCGGTACCGTCGACTCACCCTTGTTTCAGCAGGTCTG |
| TNS1 HiFi F | ggaggagaatcccggcccttCTATGAGTGTGAGCCGGAC |
| TNS1 HiFi R | agatgagtttttgttccattGATCTCTTTTGGCCGGCATTC |
